# Layer 1 NDNF Interneurons are Specialized Top-Down Master Regulators of Cortical Circuits

**DOI:** 10.1101/2023.10.02.560136

**Authors:** J Hartung, A Schroeder, Vázquez RA Péréz, RB Poorthuis, JJ Letzkus

## Abstract

Diverse types of inhibitory interneurons (INs) impart computational power and flexibility to neocortical circuits. Whereas markers for different IN types in cortical layers (L) 2-6 have been instrumental for generating a wealth of functional insights, only the recent identification of the first selective marker (NDNF) has opened comparable opportunities for INs in L1. However, at present we know very little about the connectivity of NDNF L1INs with other IN types, their input-output conversion, and the existence of potential NDNF L1IN subtypes. Here, we report pervasive inhibition of L2/3 INs (including PV- and VIPINs) by NDNF L1INs. Intersectional genetics revealed similar physiology and connectivity in the NDNF L1IN subpopulation co-expressing NPY. Finally, NDNF L1INs prominently and selectively engage in persistent firing, a physiological hallmark disconnecting their output from the current input. Collectively, our work therefore identifies NDNF L1INs as specialized master regulators of superficial neocortex according to their pervasive top-down afferents.

## Introduction

Neocortical layer 1 (L1) stands apart from all other layers in several respects: it is the thinnest of all layers and the only molecular layer, being mostly comprised of the distal dendrites of deeper pyramidal cells (PCs) and interneurons (INs) as well as extensive long-range afferents from a variety of sources^1–4^. These pathways transmit internally-generated top-down information that encodes previous experiences and current goals, and is essential for a range of higher brain functions including memory and perception^5–8^. In addition, L1 contains a small population of resident INs, which control apical PC dendrites^9^ and are thus in a prime position to control integration of top-down information in the local circuit.

While research on L1INs for a long time lagged behind the work on deeper IN types with known markers (strongly focused on parvalbumin (PV), somatostatin (SST) and vasoactive intestinal peptide (VIP) expressing populations)^10–12^, the recent identification of neuron-derived neurotrophic factor (NDNF) as a genetic marker selectively labeling approximately 2/3^rd^ of L1INs (NDNF L1INs) in both mouse and human neocortex has now opened the door for a similarly rigorous functional dissection of these cells^13–17^. To date, this work revealed that the sensory responses of NDNF L1INs are exquisitely sensitive to previous experience^13^ and arousal^15^, in line with the host of top-down afferents these cells receive, including from the higher order thalamus^7,13–15^, as well as cortical^13,15,17^ and subcortical areas^13,15,17^, such as the zona incerta^8,13,15^ and the cholinergic system^18^. These data therefore identify NDNF L1INs as central actuators of long-range top-down input in the local circuit^16^. Intriguingly, the second local type of inhibition targeting L1 is provided by SST Martinotti cells that are predominantly driven by local recurrent excitation and in consequence also display very different, often opposite *in vivo* encoding attributes^10–12,16,19–21^. Whereas both NDNF L1INs and SSTINs control distal apical PC dendrites in L1^10–14,20–22^, and SSTINs are moreover known to connect broadly to different INs types in L2-6^20,21,23^, a similarly detailed understanding of the output connectivity of NDNF L1INs is currently lacking. Since this precludes a precise definition of the circuit function of NDNF L1INs, we address these issues here. Moreover, virtually all IN types defined by a single marker gene are thought to comprise functionally distinct subtypes^24^. Following a recent landmark study^25^, we therefore validate and employ an intersectional genetic approach^26^ to selectively target the subpopulation of NDNF L1INs that coexpress neuropeptide Y (NPY). Finally, we provide a detailed characterization of the physiological attributes of NDNF L1INs, and reveal that the majority of these neurons robustly and selectively engages in persistent firing (pF), a distinct activity mode that can uncouple their action potential output from external stimulation for protracted time periods^27–31^.

## Results

### Pervasive inhibition of L2/3 IN types by NDNF L1INs

In addition to the properties and somatodendritic location of inhibition on PCs, a primary factor determining the circuit function of an IN type are its outputs to other INs. IN-IN interactions are instrumental for generating neuronal oscillations^10–12^, can gate learning and memory retrieval through disinhibition^8,32–34^, and may fundamentally add computational power to the network since IN-IN interactions are particularly pervasive in human neocortex^35^. Importantly, IN types vary widely in their IN targets and connection specificity: whereas PVINs supply little inhibition to other IN types, and VIPINs rather selectively connect to SSTINs, SSTINs themselves provide strong input to all other populations, including NDNF L1INs^9,13,23^. However, while it has been established that NDNF L1INs contact the distal apical dendrites of PCs and control dendritic calcium electrogenesis there^13–15^, their output wiring with respect to other IN types remains incompletely understood.

To define the IN-specific output wiring of NDNF L1INs and to systematically compare the properties of these connections we focused on the auditory cortex, where these cells and unidentified L1INs have been linked to learning and memory as a high-level function^1,13,16,34,36^. Employing selective Ndnf-ires-cre-ERT2 and Ndnf-ires-FlpO mouse lines we previously generated^13^ enabled us to optogenetically probe the outputs of NDNF L1INs to different L2/3 IN types in acute slices from adult animals (**Fig. 1a**, age 78-217 days, median 119). In order to obtain data under standardized conditions and to ensure comparability to previous work^13^, we calibrated the irradiance for each optogenetic activator to evoke approximately one action potential per light pulse in a train of 4 stimulations delivered at 1 Hz in NDNF L1INs using whole-cell patch-clamp recordings (**Fig. 1b,c**). To probe the connectivity from NDNF L1INs to VIPINs, we crossed Ndnf-ires-FlpO mice with VIP-cre mice^37^ and injected them with a mix of adeno-associated viral vectors (AAVs) leading to flp-dependent expression of the actuator ChrimsonR for optogenetic activation and cre-dependent expression of the fluorescent marker tdTomato for targeting of VIPINs. This revealed robust connectivity, with 10 out of 12 tested VIPINs displaying IPSCs that crossed the third standard deviation of baseline and were therefore scored as connected (**Fig. 1d-h**, see Methods for details). These IPSCs were characterized by slow rise and decay times (**Fig. 1i-k**), and displayed significant short-term depression (**Fig. 1l,m**). In conclusion, these data uncover a connection from NDNF L1INs to VIPINs that displays several of the hallmarks previously described for their inputs to PCs^13–15^.

**Figure 1.**
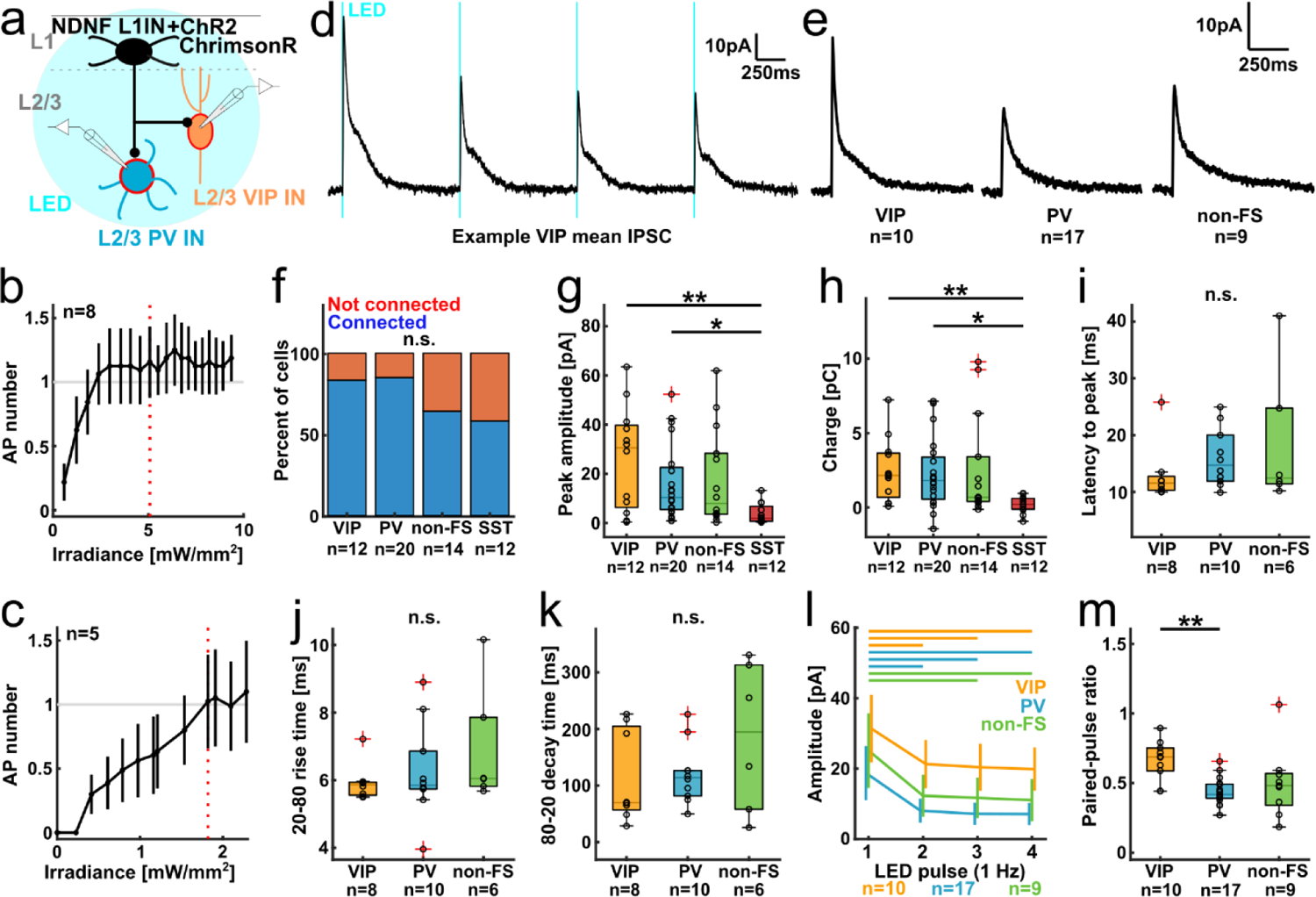
Pervasive inhibition of L2/3 IN types by NDNF L1INs. **a)** Experimental design. **b and c)** Calibration of irradiance to evoke on average one action potential per LED pulse in NDNF L1INs expressing DIO-ChR2-EYFP in Ndnf-ires-cre-ERT2 mice **(b)** or fDIO-ChrimsonR in Ndnf-ires-FlpO mice **(c)**. Red line indicates chosen irradiance. **d)** Example trace of IPSCs recorded in a L2/3 VIP IN (average of 15 trials). **e)** Grand average traces of all connected INs by type. **f)** Proportion of connected INs by type. Chi-square test p=0.26. **g)** IPSC peak amplitude. Kruskal-Wallis-Test with post-hoc Dunn’s procedure: p_group_=0.0077, p_VIP-PV_=0.98, p_VIP-nonFS_=0.87, p_VIP-SST_=0.0081, p_PV-nonFS_=1, p_PV-SST_=0.026, p_nonFS-SST_=0.13. **h)** IPSC charge. Kruskal-Wallis-Test with post-hoc Dunn’s procedure: p_group_=0.005, p_VIP-PV_=1, p_VIP-nonFS_=0.94, p_VIP-SST_=0.0085, p_PV-nonFS_=1, p_PV-SST_=0.11, p_nonFS-SST_=0.087. i-k) IPSC kinetics: latency to peak amplitude **(i)**, 20-80 rise time **(j)** and 80-20 decay time **(k)**. Kruskal-Wallis-Test: p_peak_=0.36, p_rise_=0.33, p_decay_=0.45 (Note that for analysis of IPSC kinetics, signals with an amplitude <10pA were excluded due to poor signal-to-noise-ratio). **l)** Short-term depression of NDNF L1IN elicited IPSCs. Data displayed as mean ± SEM. Bars on top indicate significant differences. Within cell type Friedman-Test with post-hoc Dunn’s procedure: VIP: p_group_=5.8e-4, p_12_=0.019, p_13_=0.033, p_14_=4e-4, p_23_=1, p_24_=0.88, p_34_=0.78. PV: p_group_=6.9e-8, p_12_=0.0086, p_13_=5.3e-6, p_14_=3e-7, p_23_=0.41, p_24_=0.14, p_34_=1. non-FS: p_group_=2.9e-4, p_12_=0.34, p_13_=0.0061, p_14_=3.5e-4, p_23_=0.61, p_24_=0.16, p_34_=0.98. **m)** Paired-pulse ratio (2^nd^/1^st^ IPSC amplitude) by cell type. Kruskal-Wallis-Test p=0.0029, post-hoc Dunn’s procedure: p_VIP-PV_=0.0021, p_VIP-nonFS_=0.08, p_PV-nonFS_=0.76.

Next, we performed analogous experiments on PVINs. For identification of these cells, we utilized recently developed viral approaches since these are compatible with the intersectional targeting of NDNF/NPY L1INs employed below. PVINs (**Fig. 1a, n**=20) were thus identified either by the PVIN-specific AAV-S5E2-tdTomato^38^, or by IN-specific AAV-mDlx-mRuby together with a maximal sustained evoked firing rate exceeding 150Hz (**Fig. S1**), a unique hallmark of these cells within cortical circuits^39^. The latter approach furthermore yielded a dataset on non-fast spiking INs (non-FS INs, n=14) for comparison. PVINs receive inputs from NDNF L1INs with high probability (**Fig. 1f**, 17/20). Whereas none of the attributes of single IPSCs differed between PVINs and VIPINs (**Fig. 1g-k**), we observed stronger short-term depression in NDNF L1IN inputs to PVINs compared to VIPINs (**Fig. 1l,m**), suggesting that inhibition of VIPINs is temporally more sustained. Together, these results therefore demonstrate that NDNF L1INs inhibit both VIPINs and PVINs as well as further, unidentified L2/3 INs, with high probability. Together with the established connectivity to other L1INs and PC distal dendrites^13–15^, this suggests that NDNF L1INs serve as master regulators of superficial neocortex.

SSTINs are a second highly connected IN type that has been suggested to function as master regulators^9,13,23^. To understand the interrelation between these two master regulators, we re-analyzed data on NDNF L1IN inputs to L2/3 SSTINs that was previously acquired in the lab using the same experimental approach and irradiance calibration strategy^13^. This revealed that NDNF L1IN inputs to SSTINs are weaker than to either VIPINs or PVINs (**Fig. 1g,h**), suggesting that the SSTIN master regulator is not strongly controlled by the NDNF L1IN master. In conclusion, these results unpack the logic of L1-led IN-IN interactions recruited by top-down afferents^1–4^, and contrast it with SSTIN-dependent circuit regulation controlled by local recurrent excitation^10–12,16,19–21^.

### NDNF L1INs comprise two physiologically distinct clusters

Given the central role of NDNF L1INs for controlling superficial circuits of neocortex, we next asked whether these cells may comprise functionally distinct subtypes, as has been shown for the general L1IN population in both mouse and human neocortex^9,16,25,40–43^. As a first step towards addressing this question, we performed whole-cell recordings on a comprehensive set of genetically-targeted NDNF L1INs (**Fig. 2a-c**, n=145), and used a standard current step protocol to exhaustively define 14 of their intrinsic physiological properties. In particular, we extracted the following parameters: action potential (AP) onset time, amplitude, threshold, half-width and maximal slope; AP afterhyperpolarization (AHP) amplitude and half-width, inter-spike interval (ISI) accommodation and variability, membrane potential, input resistance, time constant, amplitude of a depolarizing hump in the last subthreshold step, and voltage sag (**Table S1**). This revealed heterogeneity in a number of properties including AP onset time and threshold, voltage sag or resting membrane potential (Fig. 2c, **S2a**). Since all recordings were performed in current clamp, we additionally confirmed that between-cell variability in resting membrane potential does not negatively impact our measurements of the voltage sag by correlating the sag fraction measured in both current clamp and voltage clamp in the same cell in a subset of cells (**Fig. S2b-d**, Pearson R=0.94, p=4.1e-5,). Notably, a small proportion of NDNF L1INs displayed burst firing upon spike onset, and some cells showed rebound APs after hyperpolarization (**Fig. S2e,f**).

**Figure 2.**
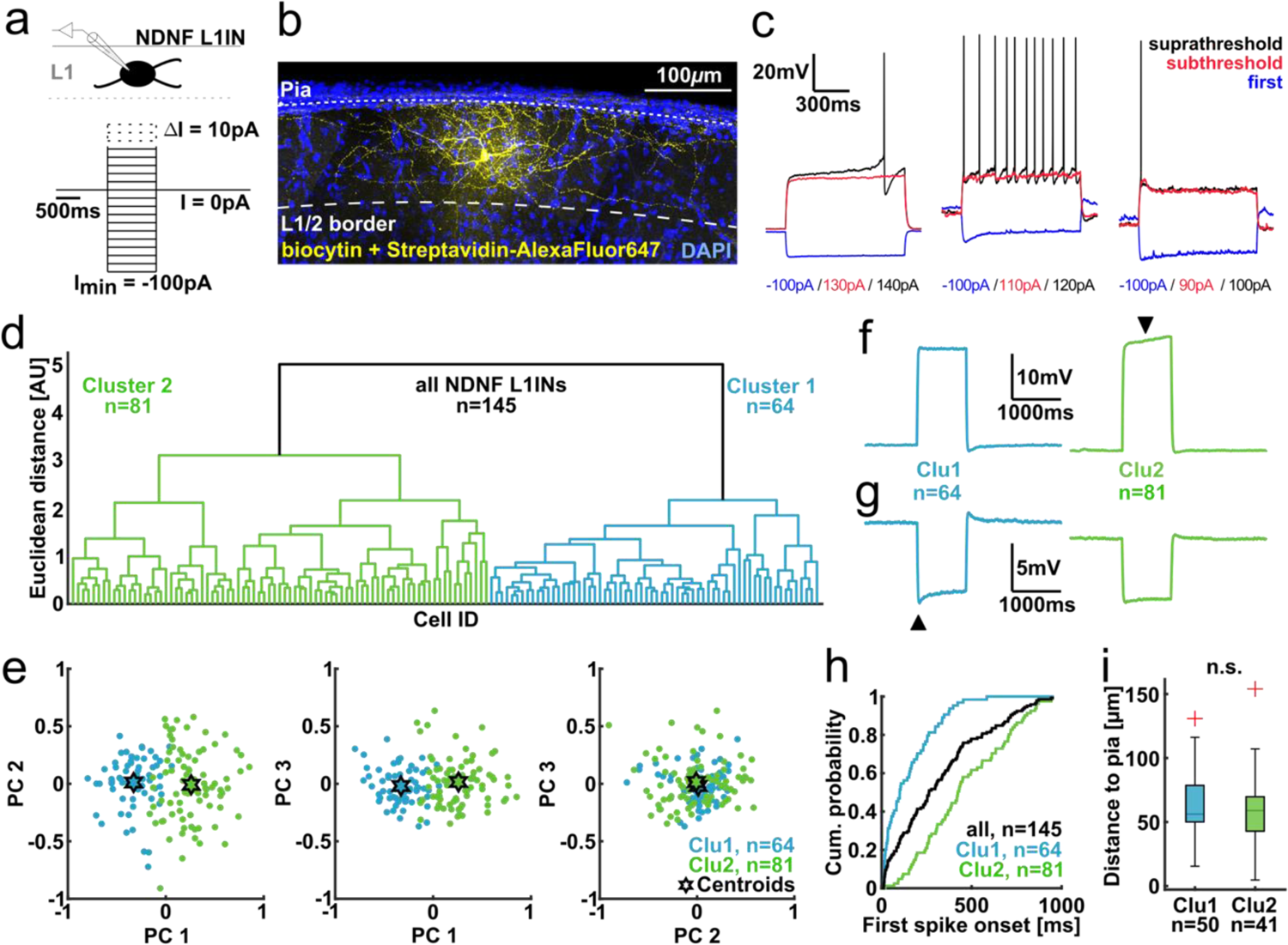
NDNF L1INs comprise two physiologically distinct clusters. **a)** Experimental design. **b)** Example histology of a recorded NDNF L1IN filled with biocytin. **c)** Example traces of the first (blue) and last subthreshold (red), and the first suprathreshold (black) step. Left: late-spiking IN with depolarizing ramp. Center: non-late-spiking IN with pronounced voltage sag in first step. Right: non-late-spiking IN with depolarizing hump in last subthreshold step. **d)** Ward’s unsupervised clustering of recorded intrinsic electrophysiological properties. Cells for which one or more intrinsic properties could not be measured were excluded. **e)** Principal component projection illustrating cluster separation. Color code corresponds to clusters obtained from **d)**. **f)** Average traces of the last subthreshold step for the 2 clusters. Black arrow indicates depolarizing ramp in cluster 2. **g)** Average traces of the step used for voltage sag measurement (black arrow). **h)** Cumulative distribution of first AP onset by cluster identity. **i)** Distance to pia of recorded NDNF INs by cluster identity. Wilcoxon rank sum test p=0.48.

To test whether these properties vary independently, or alternatively may define electrophysiological subtypes of NDNF L1INs, we subjected the data to an unsupervised clustering approach (**Fig. 2d-h**). Individual parameters were normalized in order to equilibrate their contribution (see Methods). Importantly, 4 independent approaches for evaluation of optimal cluster number converged on a local optimum of 2 main clusters for this approach (**Fig. S3a-d**), suggesting that NDNF L1INs are best described by 2 physiological clusters. Post hoc comparison revealed that the clusters differ in 9 out of 14 properties, suggesting robust separation. To further analyze the degree of cluster separation, we performed principal component analysis (PCA, Fig. 2e**, S3e**). This revealed that cluster 2 (81/145 cells) is comprised of cells in which a late AP onset typically co-occurs with higher AP thresholds, longer AP half-widths and larger AHPs. In contrast, cells in cluster 1 (64/145 cells) display early AP onsets together with larger AP amplitudes and slopes, larger voltage sags and depolarizing humps (**Fig. 2f-h, Fig. S3f-h**). Finally, we tested whether cells in the 2 clusters differ in their distance from the pia mater, but found very similar distributions (Fig. 2i). Together, these data indicate that adult auditory cortex NDNF L1INs can be described as 2 well-separated clusters based on physiology. The presence of a late-spiking phenotype with low sag fraction and depolarizing ramp that is reminiscent of previous reports from neurogliaform cells (NGFCs)^40,43,44^, and a second type characterized by early spike onset and large sag fraction and depolarizing hump is consistent with previous work both on the general L1IN population^9,16,40–43^ as well as a dissection specifically of NDNF L1INs^25^ and Lamp5-MET1INs, a class of interneurons largely overlapping with NDNF L1INs^45^.

### Intersectional genetic targeting of NDNF L1INs coexpressing NPY

The fact that NDNF L1INs comprise 2 physiologically defined clusters may suggest the presence of distinct functional subtypes^17,25^. However, it is equally compatible with a single type whose properties vary along a continuum^42,45^. To disambiguate these alternatives, further genetic identification can be used^26,45,46^. We therefore searched for an additional marker gene that may dissect the NDNF L1IN population into functionally-distinct subtypes. Given that approximately 50% of NDNF L1INs co-express neuropeptide Y (NPY)^13^, and that NPY is almost ubiquitously expressed in late-spiking, NGFCs ^44^, this suggests that NPY may be a useful additional marker. In line with this, NPY has recently been used to distinguish between NDNF L1IN subpopulations in somatosensory cortex^25^ and was found to be expressed to a varying degree within the Lmp5-MET1 IN type in L1 of visual cortex^45^.

To address this question, we choose an intersectional genetic targeting approach using Boolean cOn/fOn vectors that are only expressed if both cre and flpO recombinases are present in the same cell^26^. To this end, we crossed Ndnf-ires-FlpO mice with Npy-cre mice^47^, and injected AAV-cOn/fOn-EYFP in auditory cortex to specifically target cells that express both NDNF and NPY (Fig. 3a). To validate this approach for our system and IN type, we quantified EYFP expression and its overlap with *Ndnf* and *Npy* mRNA expression (**Fig. 3b, S4a**). Consistent with previous work, *Npy* is expressed throughout the cortical depth whereas *Ndnf* is largely confined to L1 (Fig. 3c**, S4b-d**). Importantly, EYFP expression was strongly enriched in L1 (**Fig. 3c, S4b-d**) and displayed a depth profile closely matching the *Ndnf/Npy* expression (Fig. 3c**, S4b-d**), consistent with accurate intersectional targeting. To further address this quantitatively, we profiled 2654 cells for marker overlap, and calculated sensitivity and specificity of EYFP expression (**Fig. 3c,d**; see Methods). Sensitivity quantifies the expression efficiency, and the observed sensitivity of 0.74 likely corresponds to the fact that not all cells within the injection site are transduced by the AAV. In contrast, the high specificity of 0.97 indicates a low level of aberrant expression and thus of false positives. Of note, the detection limits of the *in situ* hybridization make this a conservative estimate. In conclusion, these results demonstrate that cOn/fOn vectors can be used for accurate targeting of NDNF L1INs co-expressing the marker NPY. We note that similar experiments on the cOn/fOn AAV that would target the NDNF L1INs not expressing NPY were unsuccessful.

**Figure 3.**
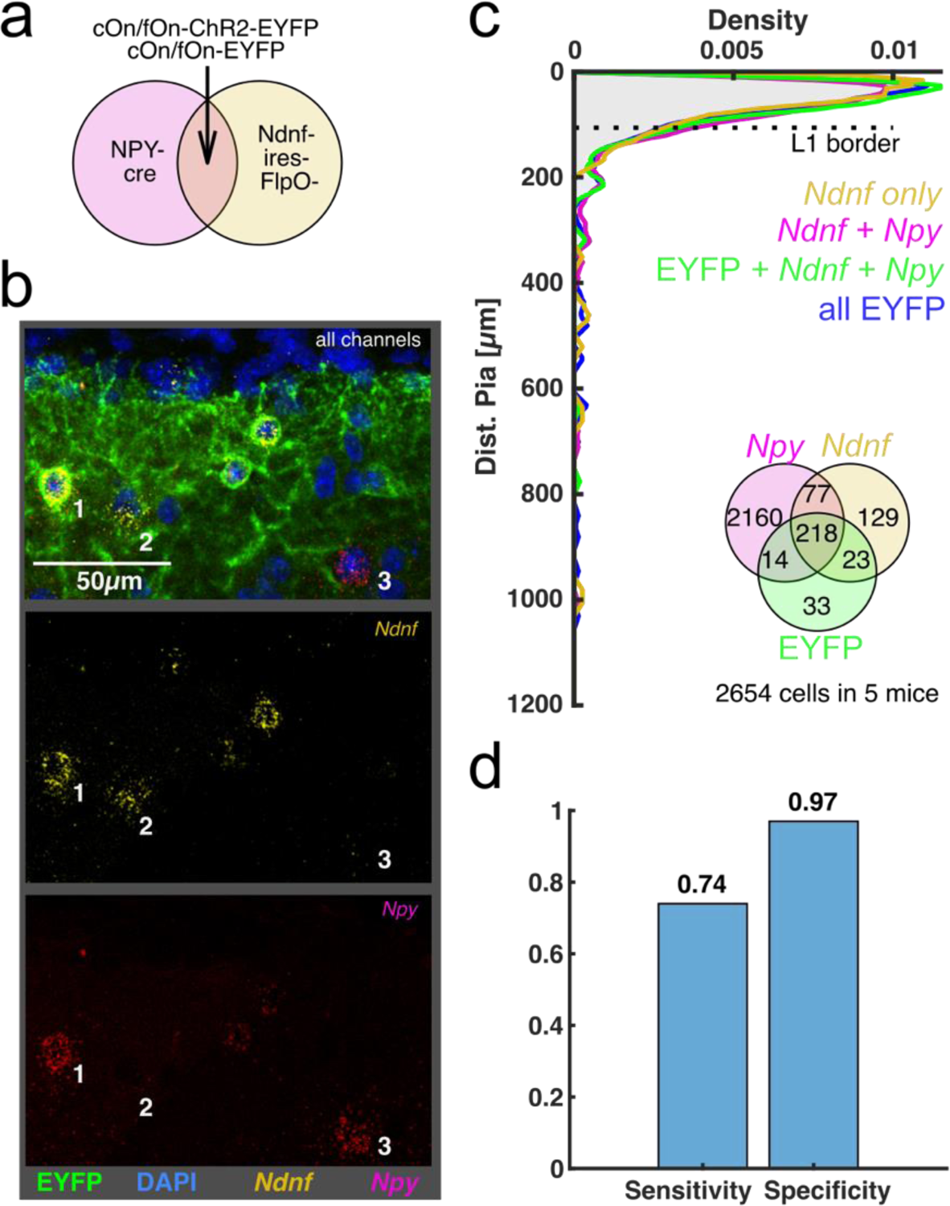
Intersectional genetic targeting of NDNF/NPY INs using a Boolean logic recombinase approach. **a)** Intersectional targeting approach. Npy-cre mice were crossed with Ndnf-ires-FlpO mice and injected with an AAV-cOn/fOn-ChR2-EYFP or AAV-cOn/fOn-EYFP, targeting the NDNF L1IN population co-expressing Npy. **b)** Example histology of intersectional EYFP (green) expression combined with *in situ* hybridization of *Ndnf* (yellow) and *Npy* (magenta) mRNA. Cell 1 co-expresses *Ndnf* and *Npy* mRNA and is EYFP positive. Cell2 is labeled for *Ndnf* but not *Npy* mRNA and is EYFP negative. Cell 3 is labeled for *Npy* but not *Ndnf* mRNA and is EYFP negative. **c)** Distribution of cells labeled for *Ndnf* but not *Npy* (yellow), cells labeled for *Ndnf* and *Npy* (magenta), cells labeled for both mRNAs and additionally expressing EYFP (green), and all cells expressing EYFP (blue). Bin width: 5µm. Inset: Venn diagram displaying number of cells labeled by *in situ* hybridization and/or expressing EYFP. **d)** Sensitivity and specificity of intersectional targeting calculated using *in situ* hybridization labeling as ground truth.

### Similar physiological attributes of NPY^+^ NDNF L1INs

Having established intersectional targeting of NDNF L1INs co-expressing NPY (NDNF/NPY L1INs), we asked whether these cells differ from the overall NDNF L1INs in their intrinsic physiological properties, and in particular, whether NDNF/NPY L1INs correspond to one of the physiological clusters we identified above (Fig. 2). To this end, we targeted these cells by whole-cell patch-clamp recordings (n=36), and recorded their attributes as in Fig. 2 (Fig. 4a). This revealed both late-spiking and non-late-spiking neurons, similar to the overall population of NDNF L1INs (Fig. 4b). Moreover, when we projected the properties of NDNF/NPY L1INs into the same principal component space defined by all intrinsic properties of NDNF L1INs (Fig. 2e), we found that the centroids are much closer to centroid of the overall NDNF L1IN population than to centroids of either cluster (Fig. 4c**, S5**). To quantitate these observations, we calculated the distance of each NDNF/NPY L1IN from the two NDNF L1IN cluster centroids. This yielded distance ratios close to zero that were statistically indistinguishable from a normal distribution (One-sample ks-test p=0.11), indicating equal distances from both NDNF L1IN cluster (Fig. 4d). Similarly, directly comparing NDNF/NPY IN distances to the two clusters in a pairwise manner revealed no significant differences (Fig. 4e). Next, we analyzed AP onset time as one of the most prominent hallmarks of the NDNF L1IN clusters (Fig. 2h), and found that NDNF/NPY L1INs closely match the overall NDNF L1IN population (Fig. 4f). Similarly, we found no significant differences between NDNF/NPY L1INs and Ndnf L1INs in 12 out of 14 of the intrinsic properties measured (**Fig. S6**). Taken together, these results indicate that NDNF/NPY L1INs do not correspond to one of the previously identified NDNF L1IN clusters, but instead most likely are identical to the overall NDNF L1IN population with regards to their intrinsic physiological attributes.

**Figure 4.**
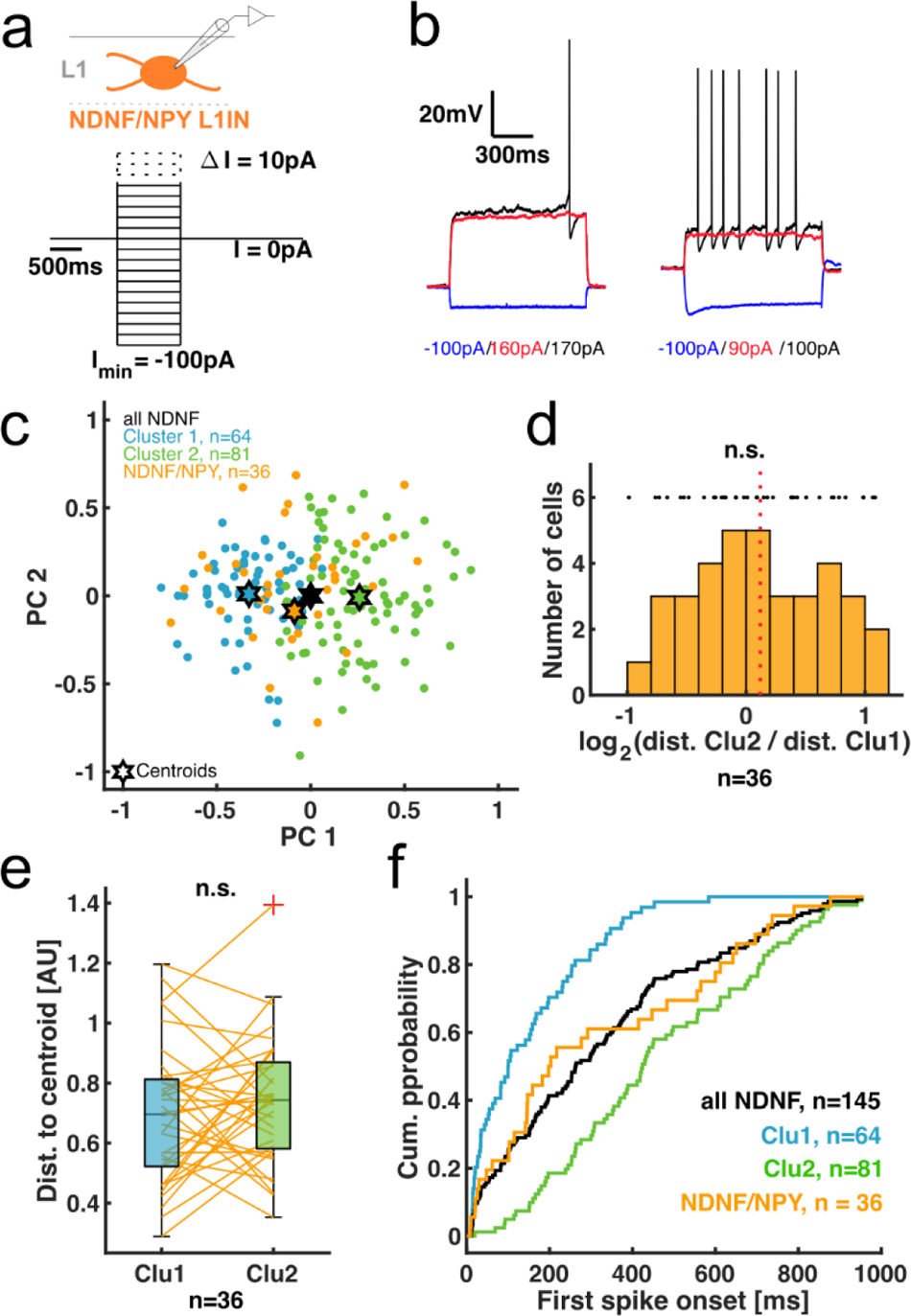
Similar physiological attributes of NPY^+^ NDNF L1INs. **a)** Experimental design. Physiological properties were measured as in Fig. 2. **b)** Example traces of the first (blue) and last subthreshold (red), and the first suprathreshold (black) step. Left: late-spiking NDNF/NPY L1IN. Right: non-late-spiking NDNF/NPY L1IN with voltage sag in first step. **c)** Projection of the intrinsic property values of NDNF/NPY cells into the PC space defined by NDNF L1IN PCA coefficients (Fig. 2e). **d)** Histogram of distance ratios of individual NDNF/NPY L1INs to centroids of NDNF L1IN clusters in full intrinsic property space. Data was log-transformed in order to scale distance ratios symmetrically around zero. 1-sample Kolmogorov-Smirnov-test p=0.11. Black dots indicate single data points, red line indicates distribution mean at 0.12. **e)** Pair-wise comparison of distances of individual NDNF/NPY L1INs from the NDNF L1IN cluster centroids in full intrinsic property space. Wilcoxon signed rank test p=0.29. **f)** Cumulative distribution of spike onset times.

### Pervasive inhibition of L2/3 IN types by NPY^+^ NDNF L1INs

Apart from their physiological attributes, NDNF/NPY L1INs may also differ from the overall NDNF L1IN population in their output connectivity as a second, functionally relevant organizational feature. To address this, we injected AAV-cOn/fOn-ChR2-EYFP into Ndnf-ires-FlpO::Npy-cre mice to be able to selectively stimulate NDNF/NPY L1INs optogenetically. Since this precludes further mouse line-based identification of their postsynaptic targets, we instead identified L2/3 INs by AAV-mDlx-mRuby and PVINs by AAV-S5E2-tdTomato, while also determining the connectivity for PCs that has not been addressed for NDNF/NPY L1INs (Fig. 5a). To ensure comparability to previous data sets, we calibrated the irradiance to evoke approximately one action potential per light pulse in a train of 4 stimulations delivered at 1 Hz (Fig. 5b). This uncovered similarly high connectivity to all neuron types tested (Fig. 5c, PCs 30/33, PV INs 21/22, non-FS INs 10/13). In contrast, IPSCs in PCs displayed larger amplitudes and charge transfer than in either PVINs or non-FS INs (**Fig. 5d-f**), in line with similar results for the overall NDNF L1IN population^13,14^. In line with our observations for NDNF L1INs (**Fig. 1i-k**), no differences in IPSC kinetics were detected between the three target cell types (**Fig. 5g-i**). Moreover, the outputs of NDNF/NPY L1INs displayed pronounced short-term depression in all three target cell types (**Fig. 5j,k**), similar to the analogous results for NDNF L1INs (**Fig. 1l,m**). In order to ensure that the observed amplitude differences between PCs and PVINs is not due to differences in AAV expression or slice quality, we performed ‘pseudo-paired recordings’^7,8^ by sequentially recording from a PC and a PVIN in close horizontal proximity and at the same cortical depth (Fig. 5l). This revealed larger IPSC amplitudes in PCs (median amplitude: PCs 31pA, PVINs 8.5pA), ruling out technical caveats as the source of the observed difference.

**Figure 5.**
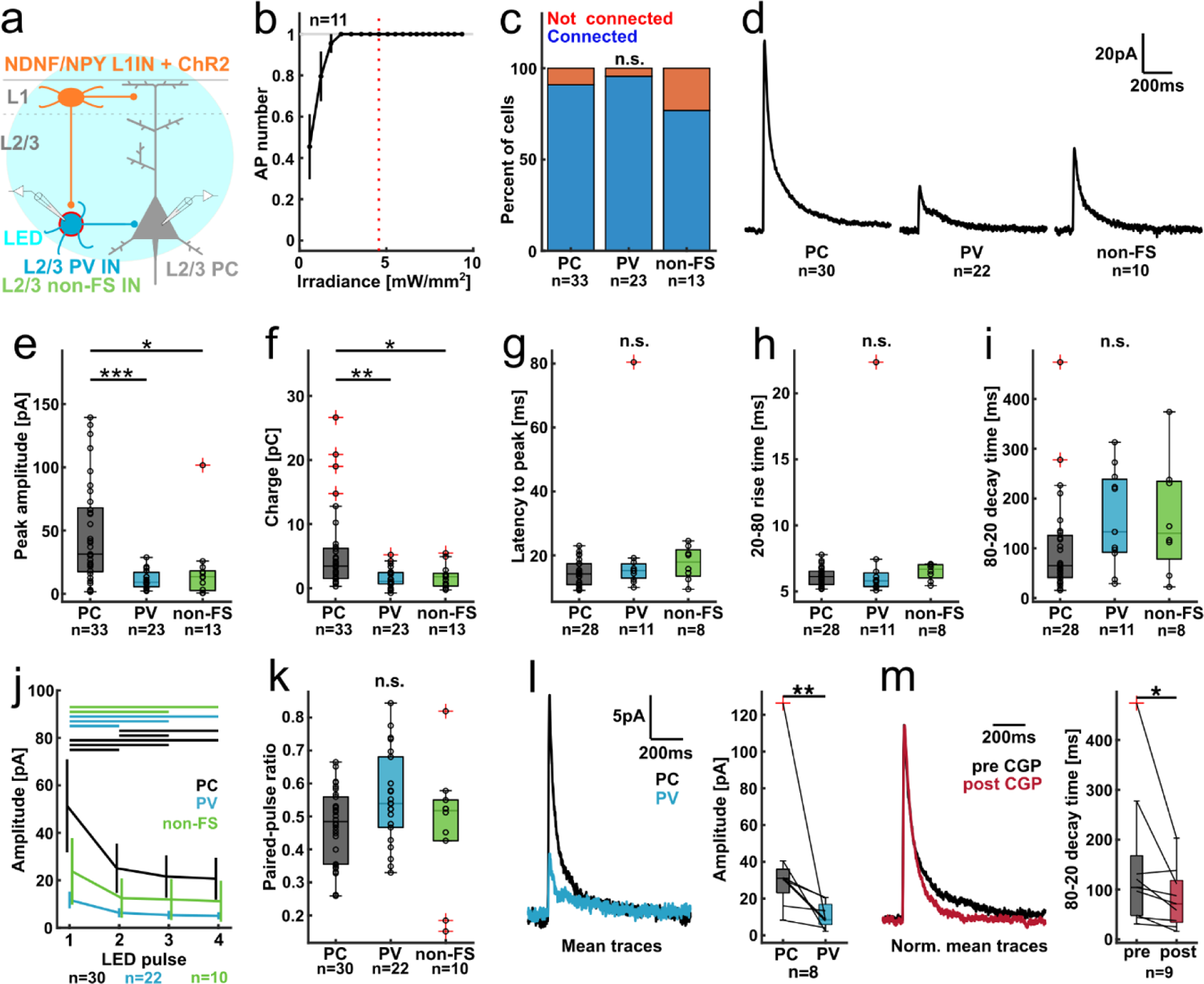
Pervasive inhibition of L2/3 IN types by NPY^+^ NDNF L1INs. **a)** cOn/fOn-ChR2-EYFP was expressed in NDNF/NPY INs in Ndnf-ires-FlpO::Npy-cre mice while in L2/3 PCs were identified morphologically, and PVINs and non-FS INs by expression of AAV-mDlx-mRuby and AAV-S5E2-tdTomato. **b)** Calibration of irradiance to evoke on average one action potential per LED pulse. Red line indicates chosen irradiance. **c)** Percentage of connected cells by cell type. Chi-square test p=0.19. **d)** Grand average traces of all connected cells by type. **e)** Peak IPSC amplitude. Kruskal-Wallis-Test with Dunn’s post-hoc procedure: p_group_=8.8e-5, p_PC-PV_=1.8e-4, p_PC-nonFS_=0.012, p_PV-nonFS_=0.96. **f)** IPSC charge. Kruskal-Wallis-Test with Dunn’s post-hoc procedure: p_group_=9.1e-4, p_PC-PV_=0.0016, p_PC-nonFS_=0.037, p_PV-nonFS_=0.98. **g-i)** IPSC kinetics: latency to peak amplitude **(g)**, 20-80 rise time **(h)** and 80-20 decay time **(i)**. Kruskal-Wallis-Test: p_peak_=0.22, p_rise_=0.2, p_decay_=0.11. **j)** Short-term depression of NDNF/NPY L1IN elicited IPSCs. Data displayed as mean ± SEM. Bars on top indicate significant differences. Within cell type Friedman-Test with post-hoc Dunn’s procedure: PC: p_group_=6.4e-16, p_12_=0.0019, p_13_=9.3e-10, p_14_=7e-15, p_23_=0.03, p_24_=6.5e-5, p_34_=0.5. PV: p_group_=2.5e-10, p_12_=0.0028, p_13_=3.2e-8, p_14_=7.6e-9, p_23_=0.11, p_24_=0.06, p_34_=1. non-FS: p_group_=8.1e-5, p_12_=0.053, p_13_=0.0041, p_14_=5.4e-4, p_23_=0.97, p_24_=0.34, p_34_=0.88. **k)** Paired-pulse ratio (2^nd^/1^st^ IPSC amplitude) by cell type. Kruskal-Wallis-Test p=0.15. **l)** Comparison of ‘pseudo-pairs’ between a PC and a PVIN. Left: Average traces. Right: IPSC amplitudes in PCs and PV INs. Wilcoxon signed rank test p=0.0078. m) Effect of CGP-mediated GABA-B receptor block in PCs. Left: Average traces. Right: IPSC 80-20 decay times pre-vs. post-CGP. Wilcoxon signed rank test p=0.02.

Finally, we addressed the contribution of GABA-B receptors as a well-described hallmark of inhibition from NDNF L1INs^13,48^. Bath application of the selective antagonist CGP-55845 (3µm) in a subset of PC recordings caused a robust reduction in IPSC decay times, indicating presence of a GABA-B receptor component (Fig. 5m, median decay time: pre CGP 104.1ms, post CGP 71.1ms), very similar to analogous data from the overall NDNF L1IN population^13^. Moreover, IPSCs recorded in PVINs also displayed a robust contribution from GABA-B receptors (**Fig. S7a,b**), suggesting that this may be a general feature of inhibition from NDNF L1INs. Taken together, our data indicate that both NDNF/NPY and NDNF INs pervasively inhibit neocortical L2/3, including PCs as well as PV and non-FS INs. Comparing inhibition by NDNF/NPY INs to that by the general NDNF INs, we found similar rates of connectivity, IPSC amplitudes and parameters, IPSC dynamics and PPR, as well as CGP effects. In combination with the similar physiology, we therefore did not observe relevant differences between these two L1IN groups.

### A specialized activity mode in NDNF L1INs

Our results so far identify NDNF L1INs as physiologically heterogeneous master regulators of superficial neocortex according to their pervasive top-down afferents^1,2,8,13–15^. We next asked whether additional hallmarks may set these cells apart from other IN types. Indeed, we frequently observed continued, autonomous action potential firing after the offset of the depolarizing current injection in NDNF L1INs (**Fig. 6a-c**, turquoise arrows). This persistent firing (pF) has been described for hippocampal and cortical NGFCs^27–30^ and is a fundamental attribute since it can effectively uncouple the activity of NDNF L1INs from ongoing synaptic input. Consistent with this being a dedicated activity mode, evoked APs and pF APs displayed clear kinetic differences, with pF APs being characterized by a clear deceleration in the upstroke (**Fig. 6d,e**). Moreover, pF episodes comprised both full APs and spikelets with lower amplitudes, whereas spikelets were not observed during evoked firing (**Fig. 6c,d, Fig. S8a**). Importantly, pF was not caused by poor cell health or recording quality since it was typically confined to a single or a few inter-step intervals, and since it completely reversed during the course of the recording (**Fig. S8b**, median ectopic AP number: pre pF 0 APs, during pF 112 APs, post pF 0 APs). Of note, in extreme cases pF could last for several minutes (**Fig. S9**).

**Figure 6.**
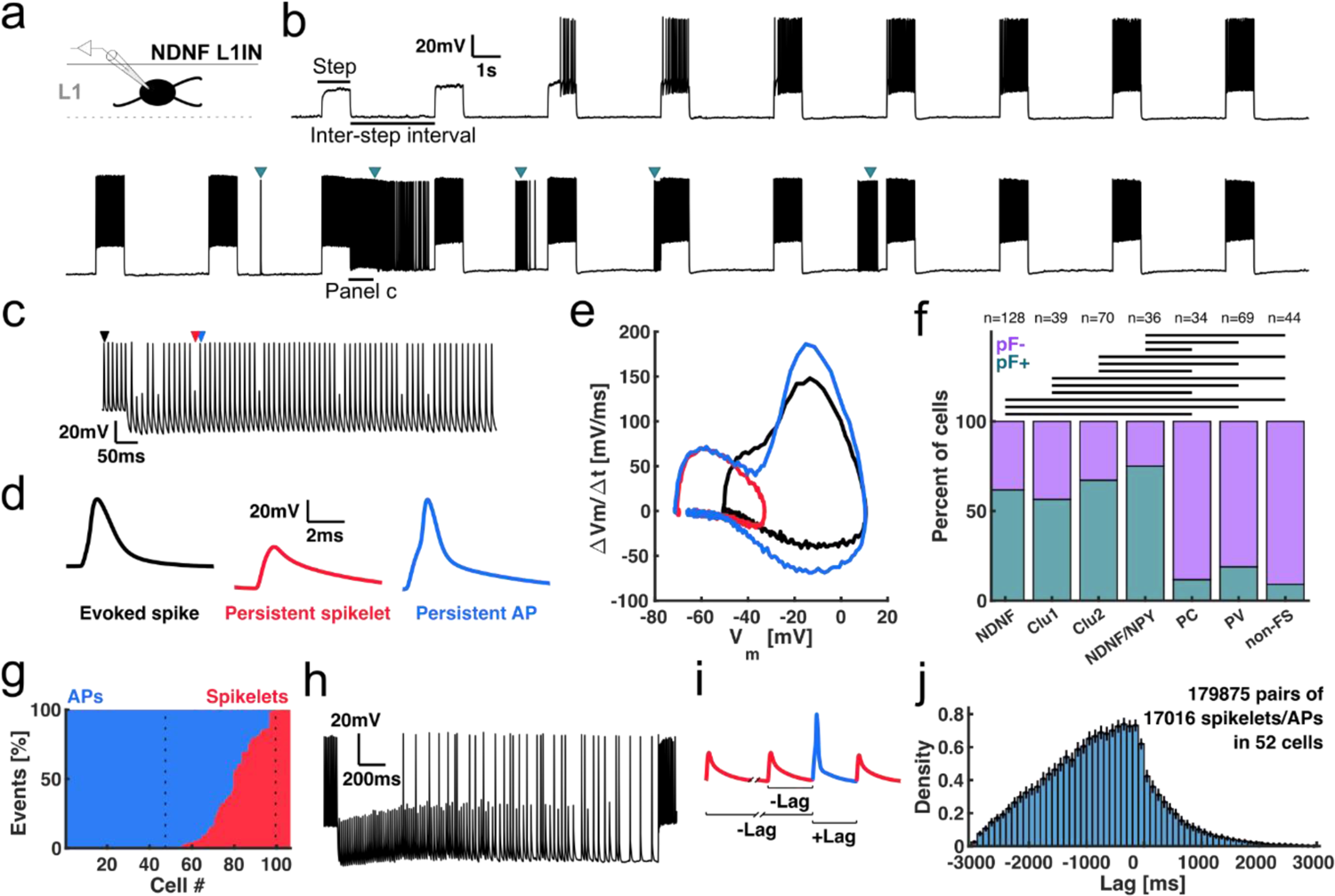
A specialized activity mode in NDNF L1INs. **a)** Experimental design. **b)** Example recording from a genetically identified NDNF L1IN. After 9 steps with current evoked spiking, the cell starts to fire spontaneously (turquoise arrows). **c)** Magnified view of pF episode in **(b)**. Colored arrows indicate events shown in (d). **d)** Waveforms of an evoked AP (black), a persistent firing spikelet (red) and a pF AP (blue). **e)** Phase plot of the events in (**d**). In contrast to the evoked AP (black), the pF AP (blue) displays a deacceleration in the AP upstroke, resulting in the prominent shoulder visible in (**d**). **f)** Occurrence of pF by cell type. 8 genetically identified VIPINs were included in non-FS INs. Chi-squared test and post-hoc Fisher’s exact test with Bonferroni correction. p_group_=3.0e-19, p_NDNF-Clu1_=1, p_NDNF-Clu2_=1, p_NDNF-NDNF/NPY_=1, p_NDNF-PC_=3.1e-6, p_NDNF-PV_=1.4e--6, p_NDNF-nonFS_=9.8e-8, p_Clu1-Clu2_=1, p_Clu1-NDNF/NPY_=1, p_Clu1-PC_=0.0016, p_Clu1-PV_=0.0021, p_Clu1-nonFS_=7.7e-4, p_Clu2-NDNF/NPY_=1, p_Clu2-PC_=1.5e-5, p_Clu2-PV_=1.8e-6, p_Clu2-nonFS_=9.9e-8, p_NDNF/NPY-PC_=61.4e-6, p_NDNF/NPY-PV_=6.6e-7, p_NDNF/NPY-nonFS_=2.6e-8, p_PC-PV_=1, p_PC-nonFS_=1, p_PV-nonFS_=1. **g)** Percentage of APs (blue) and spikelets (red) among persistent events by cell. Left dotted line indicates border between AP-only and AP-and-spikelet cells, right dotted line indicates border to spikelet-only cells. **h)** Example recording of a pF episode displaying a transition from a spikelet-only phase to a spikelet-dominated and finally an AP-dominated phase. **i)** Illustration of cross-correlation analysis. For every AP, the time lag to every spikelet was measured within all pF episodes comprising both APs and spikelets. **j)** The cross-correlation indicates that spikelets typically occur before APs during pF. Data is displayed as cross-cell mean ± SEM.

Is pF a unique capacity of NDNF L1INs? To address this, we classified a large number of recordings from different neuron types. Based on the recent report that pF can also occur as sporadic events^28^, we defined a conservative criterion of five APs occurring ectopically (without current injection) in the complete recording to score a neuron as pF (**Fig. S8c**). This revealed that approximately 2/3^rd^ of NDNF L1INs display pF (Fig. 6f, NDNF INs 79/128), whereas the proportion was much lower in PVINs (13/69), non-FS INs (4/44), and PCs (4/34). Moreover, the number of ectopic APs observed in NDNF L1INs vastly exceeded that in other neuronal types (**Fig. S8d**, median number of ectopic APs: NDNF L1INs 63 APs, PVINs: 0 APs, non-FS INs: 0 APs, PCs: 0 APs). Importantly, we observed no significant differences in either the number of cells displaying pF, nor the number of ectopic APs between NDNF L1INs and NDNF/NPY INs, nor between the two physiologically defined NDNF L1IN clusters (Fig. 6f**, S8d**), demonstrating that pF is a property of NDNF L1INs in general. In line with this, we could identify only two physiological properties within the NDNF L1IN population that differed between cells that showed and those that did not show pF (**Fig. S10**). In contrast, all other parameters, including those most strongly contributing to the variance in the NDNF L1IN population (**Fig. S3f-h**), displayed no statistically significant differences.

Within the NDNF L1IN population, both the number of inter-step intervals with pF and maximum number of ectopic APs per inter-step interval varied widely (**Fig. S8e**). Moreover, a small proportion of recorded NDNF L1INs displayed pF before the onset of evoked APs (**Fig. S8f**). Together, this indicates that the pF capacity is quite variable within the NDNF L1IN population, and that some INs are more prone to pF than others. A second parameter that notably varied is the proportion of spikelets among pF events. After defining a formal criterion to separate persistent APs and spikelets (**Fig. S8g**), we found that 44% of NDNF L1INs displayed only persistent APs, 49% showed both persistent APs and spikelets, and 7% of cells exhibited spikelets only (Fig. 6g). Moreover, in recordings that comprised both types of events, pF typically starts with spikelets and subsequently turns into a mixture which often ends with only APs (Fig. 6h). This qualitative observation was quantitated by cross-correlating APs and spikelets per inter-step interval, which confirmed that spikelets typically preceded APs across cells (**Fig. 6i,j**). In summary, we uncover pF as an attribute of NDNF L1INs that is both much less frequent and less robust in all other neuron types tested. In contrast, within the NDNF L1IN population, persistent firing is equally expressed irrespective of NPY co-expression or physiological cluster identity. We therefore conclude that this atypical activity mode represents a hallmark of the overall NDNF L1IN population.

### Persistent firing is associated with a state transition in NDNF L1INs

We next asked what biological function pF of NDNF L1INs might have. The first obvious question is whether pF is reflected in the synaptic output of these INs. We thus optogenetically stimulated NDNF L1INs at the median maximal instantaneous firing rate during pF while recording the postsynaptic response in L2/3 PCs (**Fig. 7a,b**, 50 Hz train). While this stimulation regime resulted in an initial outward current, the compound IPSC 80-20 decay time was 181.3ms, a value well within the range of decay times of IPSCs elicited by a single LED pulse by NDNF/NPY (Fig. 5i) or NDNF L1INs^13^ (**Fig. 7c, S11a**). This is consistent with the robust short-term depression of NDNF L1IN outputs even at much lower stimulation frequencies (**Fig. 1l, 5j**), and indicates that pF can only transiently cause inhibition. Given that the average duration of pF episodes by far exceeded the time window of postsynaptic transfer, we wondered whether pF might have additional, perhaps cell autonomous effects in NDNF L1INs themselves. Indeed, we noted that the onset of pF was associated with a sudden depolarization as well as the emergence of an AHP at the offset of the depolarizing current steps (**Fig. 7d-g, S11b**, step AHP). In addition, we observed that NDNF L1INs displaying pF reach higher evoked firing rates than those that do now show pF (**Fig. S11c**).

**Figure 7.**
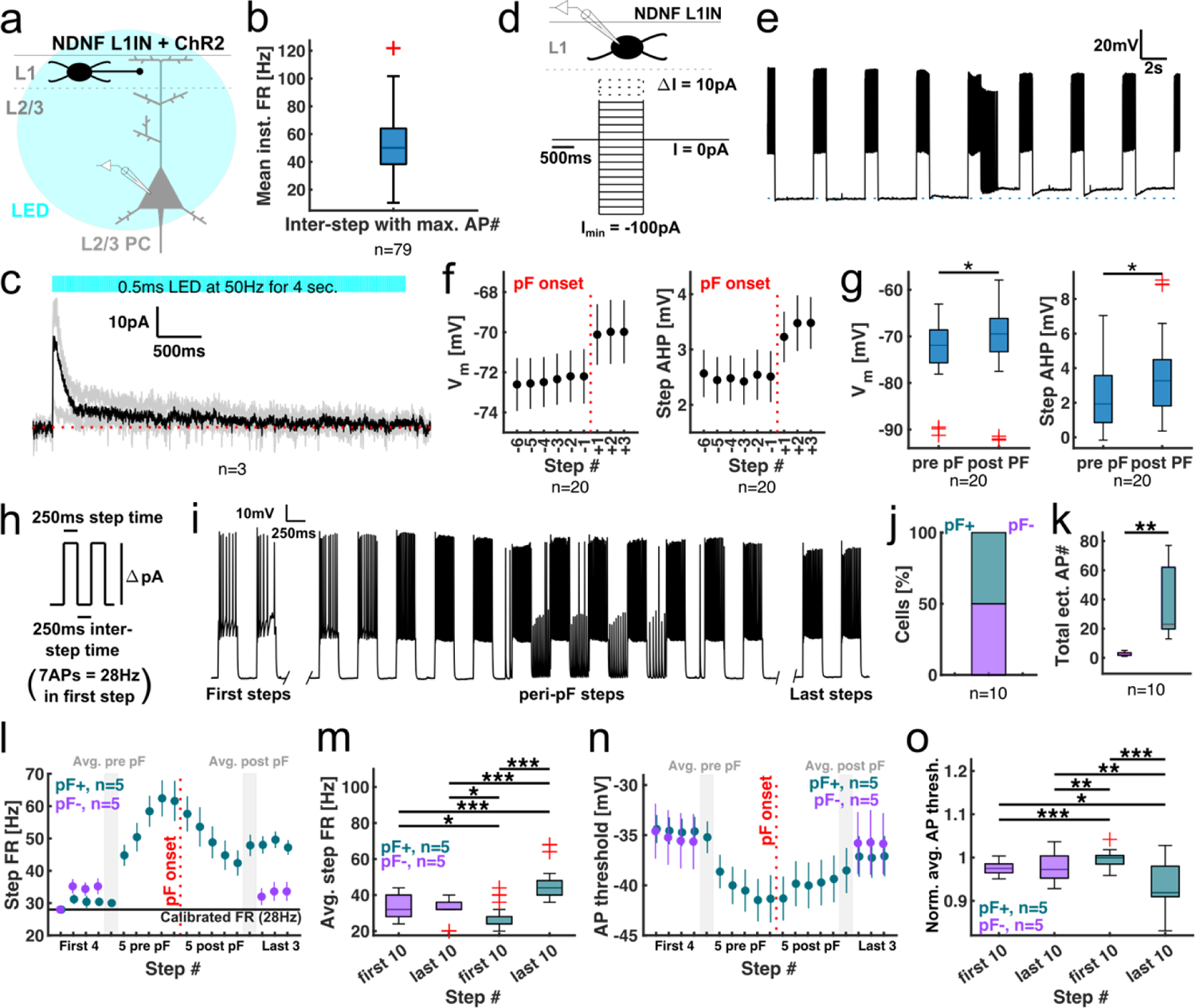
Persistent firing is associated with a state transition in NDNF L1INs. **a)** Experimental design. Stimulation was performed at 50Hz for 4s. **b)** Mean instantaneous firing rate of pF in the inter-step interval with the maximum number of ectopic action potentials. Mean = 51.2Hz / Median = 50.1Hz. **c)** Single cell (grey) and average (black) traces of IPSCs in L2/3 PCs. Red line indicates pre-IPSC baseline. **d)** Experimental design. **e)** Example recording highlighting changes in resting membrane potential and step AHP after onset of persistent firing. **f)** Left: Changes in resting membrane potential pre vs. post pF onset. Right: Changes in step AHP pre vs. post pF onset. Displayed as mean ± SEM. Red line indicates onset of pF. All recordings of NDNF L1INs included in which there was no further pF for three inter-step intervals after pF onset. **g)** Comparison of the 3 steps preceding vs. the 3 steps following pF onset. Left: Resting membrane potential. Wilcoxon rank sum test p=0.02. Right: Step after-hyperpolarization. Wilcoxon rank sum test p=0.15. **h)** NDNF L1INs were stimulated with constant amplitude current steps of 250ms duration at 2Hz. Amplitudes were calibrated for each cell to elicit APs at 28Hz in the first step. **i)** Example recording of a NDNF L1IN displaying pF. Left: First two steps. Center: peri-pF steps. Right: Last two steps in recording. **j)** Percentage of NDNF L1INs displaying pF during the protocol. **k)** Total number of ectopic APs in cells that do or do not display pF. Wilcoxon rank sum test p=0.0079. **l)** Evoked firing rates over time. Data displayed as mean ± SEM. Red line indicates onset of pF. **m)** Comparison of firing rates in first 10 vs. last 10 steps by presence or absence of pF. Kruskal-Wallis-test p_group_=3e-20. Post hoc Dunn’s test p_first--last-_=1, p_first--first+_=0.013, p_first--last+_=1.9e-9, p_last--first+_=0.035, p_last--last+_=2.3e-10, p_first+-last+_=0. **n)** Same as (**l**) for the threshold of evoked APs. Data displayed as mean ± SEM. Red line indicates onset of persistent firing. **o)** Same as (**m**) for the threshold of evoked APs. Kruskal-Wallis-test p_group_=2.5e-10. Post hoc Dunn’s test p_first--last-_=0.98, p_first--first+_=3.8e-4, p_first--last+_=0.025, p_last--first+_=0.0052, p_last--last+_=0.0025, p_first+-last+_=4.1e-11.

Since the recording protocol with escalating current steps makes it hard to disambiguate the effects of pF from those of stronger exogenous stimulation, we switched to a series of current steps with constant amplitude calibrated to evoke APs at 28Hz, a frequency that provides enough dynamic range to detect changes in firing rate in either direction, in the first step (**Fig. 7h,i, S11d**). Consistent with our previous results, when applying the same criterion for pF as above, a large fraction of recorded NDNF L1INs displayed pF (**Fig. 7j,k, S11d,e**). Intriguingly, in the cells displaying pF, we observed that evoked firing rates started to increase a few steps before, and peaked shortly before, pF onset (**Fig. 7l,m, S11f**). Evoked firing rates decreased again after pF onset but remained constantly elevated relative to pre-pF over tens of seconds until the end of the recording (**Fig. 7l,m, S11f**). In contrast, these dynamics were completely absent in NDNF L1INs without pF. In fact, except for an initial, much smaller increase after the first step, firing rates were practically constant. Strikingly, we observed similar dynamics for the threshold of evoked APs, which started to decrease towards pF onset, with a minimum immediately before pF onset (**Fig. 7n,o, S11g**). Similar to evoked firing rates, AP thresholds started to increase again after pF onset, but did not return to pre-pF values until the end of the recording. Again, NDNF L1INs without pF showed no such dynamics. Notably, this state of enhanced excitability was long-lasting and in particular the increased firing rates could be sustained for as long as the cell was continuously stimulated (up to several minutes). In contrast, a prolonged resting period without any stimulation (at least ∼30 seconds) resulted in a return to the pre-pF electrophysiological state. This cycle of stimulation, persistent firing and transition to a highly excitable state, and return to the initial state after a period of rest could be repeated many times in the same cell. Together, these data indicate that whereas pF only causes transient postsynaptic inhibition, it is associated with a profound and long-lasting transition in the intrinsic state of NDNF L1INs towards greater excitability.

## Discussion

NDNF L1INs have recently emerged as a type of IN in mouse and human neocortex that is distinct from the much more intensely investigated populations expressing PV, SST and VIP^13–17,43^. Despite the fact that NDNF L1INs display a range of specialized and intriguing attributes, including robust encoding of experience^13^ and arousal^15^, rapid neuromodulation^43^ and input from a large number of brain-wide afferents conveying top-down information^1,8,13–17^, our understanding of the intrinsic and circuit function of these neurons has remained incomplete. Here, we use *in vitro* patch-clamp recordings combined with single and intersectional genetic targeting, *in situ* hybridization, unbiased clustering and optogenetics to close this gap in our understanding.

### Integration into the neocortical circuit

Our data reveal that NDNF L1INs not only inhibit the distal apical dendrites of PCs in L1^13,14^, but at the same time pervasively inhibit the majority of L2/3INs, including those expressing VIP and PV. As NDNF L1INs themselves receive strong and diverse long-range top-down projections^7,8,13–15,17^, these findings therefore define NDNF L1INs as top-down master regulators of the superficial cortex^9^ (Fig. 8). Intriguingly, the one IN type that is not substantially inhibited by NDNF L1INs are SSTINs^13–15^. Like NDNF L1INs, they have been described as cortical master regulators^9^ based on the fact that they, too, broadly inhibit cortical INs^9,23^ as well as PC apical dendrites^20,22^. However, SSTINs are most strongly driven by local recurrent activity^19–21^, and moreover strongly inhibit NDNF L1INs^13^ (Fig. 8). In summary, this similarity in output connectivity together with the contrast in *in vivo* encoding^13–17,19–21^, potentially opposing roles during learning^13,16^ and the largely unidirectional inhibition between the two cell types^9,13–15^ define the working hypothesis of two hierarchically organized cortical master regulators recruited by local recurrent versus long-range top-down information. Future experimental and theoretical work addressing how these IN types interact *in vivo* during integration of bottom-up and top-down information is therefore required^13,16,49^.

**Figure 8:**
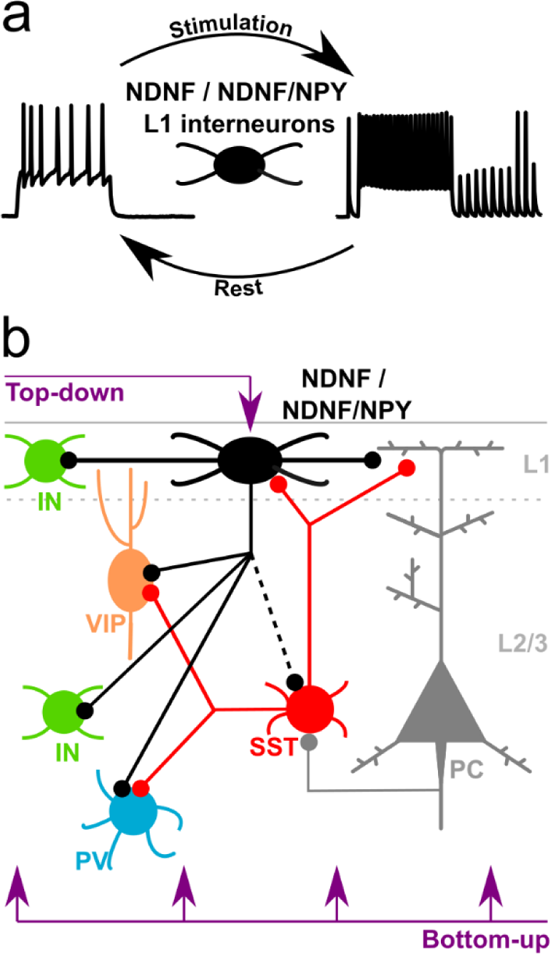
NDNF INs are master regulators of superficial cortical circuits that undergo an activity-dependent transition of electrophysiological state. **a)** After sufficient recruitment, the majority of NDNF L1INs undergo a state change triggering pF in the absence of further stimulation together with enhanced excitability. This state is maintained by continued stimulation, and reverts after a period of rest. **b)** NDNF L1INs are top-down master regulators of superficial cortical circuits, controlling L2/3 PCs, PVINs, VIPINs and unidentified L1^13^ and L2/3 INs. Previous work suggests that SSTINs represent a second master regulator driven predominantly by local recurrent excitation. Connectivity between the two masters is largely unidirectional, suggesting that SSTINs may be able to override the NDNF L1IN master regulator.

Similarly, connectivity between NDNF L1INs and superficial VIPINs is unidirectional as well: while we find that NDNF L1INs inhibit superficial VIPINs, the reverse is not the case^13^. Both IN types are targeted by top-down afferents^13–15,21,50^ and key elements in local disinhibitory circuits^14,15,21,23,50^. However, whereas NDNF L1INs directly inhibit PC dendrites and indirectly – via PVINs – disinhibit PC somata, VIPINs indirectly – via SSTINs – disinhibit PC dendrites and – via disinhibition of PVINs – enhance inhibition of PC somata^13–15,20,21,23,50^. The two IN types therefore produce opposing effects on PCs, and the unidirectional connectivity we uncover suggests a functional hierarchy between the two disinhibitory circuits.

### Potential subtypes of NDNF L1INs

Given the pervasive finding that IN types identified by single genetic markers can be further subdivided into distinct subclasses^24^, we utilized intersectional genetic targeting to address this issue for NDNF L1INs. While the overall population of NDNF cells falls apart into two physiological clusters, we found no correspondence of these attributes with co-expression of NPY. Moreover, NPY ^+^ and NPY^-^ NDNF L1INs display similar incidence of pF, IPSC kinetics and circuit connectivity, together indicating that there is insufficient functional distinction for dubbing NPY^+^ NDNF L1INs a distinct subclass of IN. In contrast, previous work in juvenile somatosensory cortex (age P21-P53)^25^ reported a clear separation between NPY ^+^ and NPY^-^ NDNF L1INs, while a continuum of attributes between NPY ^+^ and NPY^-^ L1INs expressing Lamp5-MET1 (a population largely overlapping with NDNF in L1^45^) was found in visual cortex of young adults (age P45-P70)^45^. While these apparent discrepancies may thus also be due to area differences, a perhaps more likely explanation relates to the fact that, different from most canonical IN markers, expression of NPY in INs expands continuously with age^51^. We therefore submit that NPY may be a good marker to define NDNF L1IN subpopulations in juvenile neocortex, but becomes less specific with age. Moreover, hippocampal NGFCs can undergo a form of activity-dependent plasticity essentially transforming a late-spiking phenotype with depolarizing ramps and high AP thresholds into a non-late-spiking phenotype with depolarizing humps and low AP thresholds^27^. These differences map precisely onto the NDNF L1 IN clusters we obtained, indicating that they might correspond to different electrophysiological states rather than to hard wired IN subtypes.

### Persistent firing, a specialized activity mode of NDNF L1INs

Whereas the overwhelming number of neuron types exclusively functions as *signal integrators*, our data uncover that after sufficient stimulation, the majority of NDNF L1INs switches to becoming *signal generators* by persistently producing APs in the absence of stimulation (Fig. 8). Persistent firing originates in the distal axon^28,29^ and has primarily been observed in hippocampal and cortical NGFCs^27–30,52^. In line, we find that while pF occurs in a large proportion of NDNF L1 INs, we observe it only rarely and much more weakly in other cortical cell types.

Functionally, our results indicate that pF does not elicit sustained postsynaptic inhibition due to the robust short-term depression we observe. In contrast, we find that pF is associated with a profound and long-lasting transition in NDNF L1IN electrophysiological state, including changes in resting membrane potential, step AHP, action potential threshold and firing rate. Our data therefore argue for a cell-autonomous effect of pF, that may for instance modulate the induction of long-term synaptic plasticity at the afferents of these cells^27,53^. In the temporal domain, the long-lasting nature of pF is in line with several other NDNF L1IN characteristics that occur on protracted timescales, including late-spiking, slow single IPSC kinetics, pronounced short-term depression and the fact that top-down inputs often display much greater latencies than bottom-up signals^1^. Together, this argues for NDNF L1INs as a neuron type that is optimized for slow signals.

## Methods

### Animal subjects

All mouse lines used were maintained on a C57BL6/J background. Mice were housed under a 12h light/dark cycle and provided with food and water *ad libitum*. After surgical procedures, mice were individually housed. All animal procedures were executed in accordance with institutional guidelines, and approved by the prescribed authorities (Regierungspräsidium Darmstadt and Freiburg).

### Surgery

Mice were anesthetized with isoflurane (induction: 4%, maintenance: 1.5-2%) in oxygen-enriched air (Oxymat 3, Weinmann, Hamburg, Germany) and fixed in a stereotaxic frame (Kopf Instruments, Tujunga, USA). Core body temperature was maintained at 37.5C via a feed-back controlled heating pad (FHC, Bowdoinham, ME, USA). Analgesia was provided by local injection of ropivacain under the scalp (16.7mg/kg, Ropivacain-HCl B.Braun) and subcutaneous injection for systemic action of metamizol (200mg/kg, Novaminsulfon-ratiopharm) and meloxicam (1-2mg/kg, Metacam Boehringer Ingelheim). Adeno-associated viral vectors (AAV, serotype 2/1, 2/5 or 2/9, max. 700nl) were injected from glass pipettes connected to a pressure ejection system (PDES-02DELA-2, NPI, Germany) into auditory cortex at the following coordinates: 2.46 mm posterior from bregma, 4.6 mm lateral of midline, depth below cortical surface 100-600µm.

### Slice preparation for electrophysiology experiments

Mice of both sexes (age 78-217 days, median 119) were deeply anaesthetized with isoflurane (5%) in oxygen-enriched air (Oxymat 3, Weinmann, Hamburg, Germany), and decapitated into carbonated, ice cold slicing solution. A Leica VT 1200S vibratome was used to obtain 350µm thick coronal slices from auditory cortex. Slices were directly transferred to carbogenated slicing solution at 33°C for 10min, and then further transferred to carbogenated standard ACSF at room temperature. After 30-60 minutes recovery time, slices were used in whole-cell patch-clamp experiments. Slicing solution contained (in mM) 93 NMDG, 93 HCl, 2.5 KCl, 1.2 NaH2PO4, 30 NaHCO3, 20 HEPES, 25 glucose, 5 sodium ascorbate, 2 thiourea, 3 sodium pyruvate, 10 MgSO4 and 0.5 CaCl2 and was calibrated to a pH of 7.3-7.4 and an osmolarity of 300-310mOsm. Standard ACSF contained (in mM) 125 NaCl, 3 KCl, 1.25 NaH2PO4, 26 NaHCO3, 10 glucose, 1 MgCl2 and 2 CaCl2 and was calibrated to an osmolarity of 300-310mOsm.

### Patch clamp electrophysiology

Slices were held in a recording chamber at 33°C and perfused with ACSF (2-4 mL/min). Cells were visualized for patching using differential interference contrast microscopy (Scientifica) or under epifluorescence for identification using an LED (488 or 565 nm, Cool LED) with a water immersion objective (Olympus LUMPlanFLN40xW) and a CCD camera (Scientifica SciCam Pro). Cells were recorded in whole-cell patch-clamp recordings using pipettes pulled from standard-wall borosilicate capillaries using a (3.5-6 MOhm, DMZ Zeitz-Puller). Intracellular solution contained (in mM): 140 K-gluconate, 10 KCl, 10 HEPES, 4 Naphosphocreatine, 4 ATP-Mg, 0.4 GTP and biocytin (4 mg/mL) and was calibrated to pH 7.3 with KOH and an osmolality of 290-300 mOsm. A Multiclamp 700B amplifiers (Axon Instruments, CA) was used for whole-cell voltage clamp or current clamp recordings, together with a Digidata1550 (Molecular Devices) for digitization. Recordings were low pass filtered at a 10kHz using a Bessel filter and digitized at 50kHz. Series resistance was routinely compensated in voltage clamp and recordings were excluded if access resistance exceeded 30MOhm.

### Intrinsic electrophysiological properties

NDNF L1INs were identified genetically by expressing AAV2/5.EF1a.DIO.hChR2(H134R)-EYFP.WPRE.hGH (PennVector Core) in Ndnf-ires-cre-ERT2 animals (B6(Cg)-*Ndnf^tm^*^1.1^(cre/ERT2)*^Ispgl^*/J, The Jackson Laboratory), by expressing AAV2/1-EF1a-fDIO-EYFP-WPRE (UNC Vector Core) in Ndnf-ires-FlpO animals (B6(Cg)-*Ndnf^tm^*^1.1^(flpo)*^Ispgl^*/J, The Jackson Laboratory) or by crossing Ndnf-ires-cre-ERT2 animals with an ai9 tdTomato reporter line (B6.Cg-*Gt(ROSA)26Sor^tm9(CAG-tdTomato)Hze^*/J, The Jackson Laboratory). Intersectional NDNF/NPY L1INs were identified by crossing Ndnf-ires-FlpO mice with Npy-cre (B6.Cg-*Npy^tm1(cre)Zman^*/J, The Jackson Laboratory) mice and expressing AAV2/5-hSyn-Con/Fon EYFP-WPRE (UNC Vector Core) or AAV2/5-hSyn-Con/Fon-hChR2(H134R)-EYFP-WPRE (UNC Vector Core). Intrinsic properties were recorded in current clamp at the cells spontaneous resting membrane potential using a protocol consisting of 1 second baseline without any current injection, a 1 second current injection step, and 2 seconds baseline. Current steps started at −100pA or −40pA and increased by 10pA every sweep. Cells were included for analysis if they met general recording quality criteria and all intrinsic properties could be estimated.

### Connectivity

For optogenetic probing of connectivity from NDNF L1INs to VIPINs, Ndnf-ires-FlpO mice were crossed with VIP-cre mice (STOCK *Vip^tm1(cre)Zjh^*/J, The Jackson Laboratory). AAV2/1-EF1a-fDIO-ChrimsonR-wpre-sv40 (Vector Biolabs) and AAV2/1-EF1a-fDIO-EYFP-WPRE (UNC Vector Core) were used for identification and optogenetic activation of NDNF L1INs, and AAV2/1.CAG.Flex.tdTomato.WPRE.bGH (PennVector Core) was used for identification of VIPINs. For probing connectivity from NDNF L1INs to PCs and PV and non-FS INs, AAV2/5.EF1a.DIO.hChR2(H134R)-EYFP.WPRE.hGH (PennVector Core) was used in Ndnf-ires-cre-ERT2 animals for identification and optogenetic activation of NDNF L1INs, and pAAV-mDLX-NLS-mRuby (addgene) or pAAV-S5E2-Gq-P2A-dTomato (addgene) were used to aid identification of INs in general for PVINs in particular. Pyramidal cells were identified morphologically. Interneurons identified by pAAV-mDLX-NLS-mRuby with a maximum evoked firing rate >150Hz were counted as PVINs. For probing connectivity from NDNF/NPY L1INs to L2/3INs, Ndnf-ires-FlpO animals were crossed with Npy-cre animals and AAV2/5-hSyn-Con/Fon-hChR2(H134R)-EYFP-WPRE was used to identify and optogenetically activate NDNF/NPY L1INs. L2/3 pyramidal cells and PV and non-FS INs were identified in the same manner as in Ndnf-ires-cre-ERT2 animals described above. Calibration experiments were conducted for all optogenetic activators used in order to determine a LED power eliciting approximately one action potential per 0.5ms LED pulse in the NDNF or NDNF/NPY L1IN population, by recording NDNF or NDNF/NPY L1INs in current clamp and applying the same protocol as for actual connectivity recordings (see below). The calibrated LED power was used for all further experiments using a given optogentic activator. Connectivity was probed in VC with recorded cells clamped to −50mV and delivering a train of four 0.5ms LED pulses at 1Hz for 10-15 sweeps. Recordings with a negative holding current at −50mV were excluded as we assumed poor cell health. Light was delivered in full-field application through the objective, using a LED (Cool LED) at 488nm (for ChR2) or 565nm (for ChrimsonR). For offline processing, signals were averaged across sweeps and filtered with a 4-pole Butterworth filter with a Fc of 500Hz. Cells were considered connected if the average signal after the first LED pulse exceeded the third standard deviation of the pre-LED pulse baseline. Only the first IPSC in every train was used for analysis of IPSC characteristics. IPSCs with amplitudes <10pA were excluded from the analysis of IPSC kinetics due to low SNR. Amplitude was calculated as the difference between the pre-LED baseline and the maximum value of an IPSC. Charge was calculated as the integral over one second after the LED stimulus. Latency to peak was calculated as time between LED onset and IPSC maximum value. Rise time was calculated as times between the points at which the signal crossed 20% and 80% of the amplitude on the signal upstroke. Decay time was calculated as time between the points at which the signal crossed 80% and 20% of the amplitude on the signal downstroke. GABA-B contributions were measured by application of 3µm CGP-55845 and comparison of pre and post CGP application IPSC decay times. Recordings were excluded it access resistance differed by more than 20% between pre and post CGP recordings.

### Persistent firing

Persistent firing in NDNF L1INs was investigated using the same recordings as for intrinsic electrophysiological properties. Persistent firing was also investigated in NDNF/NPY L1INs as well as all L2/3 cell types included in analysis of connectivity, following the same protocol as in NDNF L1INs (although in a fraction of cells, current step amplitude increased by 20pA with every sweep). For all recordings using the 1-second current step protocol, only recordings were considered for analysis of persistent firing if cells reached depolarization block at the end of the recording, in order to prevent biasing the rates of occurrence by recordings which might have been terminated too early. In addition, in genetically identified NDNF L1INs, persistent firing was investigated using a protocol consisting of 250ms current injections steps at 2Hz, with 19 steps per sweeps and 10 sweeps. In this protocol, injected current amplitudes were constant and calibrated to elicit action potentials at 28Hz during the first step.

### Fluorescent in situ hybridization (FISH) with immunohistochemistry

Ndnf-FlpO::NPY-Cre mice were injected with AAV2/5-hSyn-Con/Fon-EYFP-WPRE (UNC) in ACx. 6 weeks post-injection, mice were anesthetized with isoflurane, sacrificed and the brains were then dissected, embedded in Tissue-Tek O.C.T. compound (Sakura) and frozen in isopentane at −55 to −60°C. 16 μm-thick sections from these fresh frozen brains were prepared using a cryostat (Leica) and mounted on SuperFrost Plus microscope slides (Thermo Scientific). Sections were screened for fluorescence in ACx (Zeiss Axio Zoom) and then stored at −80°C until FISH was performed using the RNAscope Fluorescent Multiplex Reagent Kit (#320850, Advanced Cell Diagnostics). Heating steps were performed using the HybEZTM oven (Advanced Cell Diagnostics). Tissue sections were treated with pretreatment solutions and then incubated with RNAscope probes (Mm-Npy-C2, #313321-C2; Mm-Ndnf-C3, #447471-C3) diluted in probe diluent (#300041), followed by amplifying hybridization processes. This was followed by immunofluorescent staining of EYFP. Blocking solution consisted of 10% normal goat serum and 0.5% triton in PBS-0.2% gelatin. Primary antibody chicken anti-GFP (Aves) was diluted in 5% normal goat serum and 0.5% triton in PBS-0.2% gelatin. DAPI was used as a nuclear stain. Prolong Gold Antifade (Thermo Scientific) was used to mount slides. Images were acquired on a confocal microscope (Zeiss LSM 880). EYFP-expressing cells, their distance from the pia and overlap with markers were quantified using a custom written MATLAB (MathWorks) script. The sensitivity is the probability to detect a signal (EYFP) when there actually is a signal, and was calculated as number of true positives / (number of true positives + number of false negatives). True positives are cells labeled by EYFP and both, *Ndnf* and *Npy* mRNA. False negatives are cells that are labeled by both mRNAs but not by EYFP. The specificity is the probability not to detect a signal when there is no signal, and was calculated as number of true negatives / (number of true negatives + number of false positives). True negatives are cells not labeled by EYFP and labeled by only one, *Ndnf* or *Npy* mRNA. False positives are cells labeled by EYFP and maximally one mRNA.

### Data analysis and statistics

All statistical analyses were done using MATLAB. For clustering analyses, each intrinsic property from NDNF and NDNF/NPY L1INs was normalized to a range between 0 and 1. Data from NDNF and NDNF/NPY L1INs was normalized together to ensure comparability. For unsupervised clustering, Ward’s method was used using normalized data from NDNF L1INs. PCA was calculated using normalized data from NDNF L1INs. Comparison with data from NDNF/NPY L1INs was done by projecting normalized NDNF/NPY L1IN intrinsic properties into PC space using PCA coefficients obtained from calculating PCA on NDNF L1IN data. All data was tested for normality before statistical testing using the one-sample Kolmogorov-Smirnov test, a Lilliefors test, Jarque-Bera test and Anderson-Darling test. If all test indicated a normal distribution, parametric methods were used, and non-parametric methods otherwise. In addition, data sets containing both normal and non-normal distributed data were analyzed using only non-parametric methods for comparability. For parametric test, a paired t-test was used for paired comparisons with two samples. For non-parametric tests, the Kruskal-Wallis test was used for group-wise between group comparisons, and the Friedman test for group-wise within-group comparisons. If they proved significant at alpha of 0.05, they were followed by the Dunn-Šidák procedure for pairwise testing and multiple test correction. Non-parametric, non-paired comparisons between two samples were done using Wilcoxon’s rank sum test. Non-parametric, paired comparisons between two samples were done using Wilcoxon’s singed rank test. Statistics indicated in plots read as: n.s. p >= 0.05, * p <0.05, ** p<0.01, *** p < 0.001.

## Data availability

All source data will be deposited at zenodo.org, and will become publicly available as of the date of publication. Requests for materials should be directed to JJL. All custom scripts used for data analysis and statistics will be made available on GitHub upon publication. Requests related to code should be directed at JH.

## Author contributions

JH and JJL conceived the project. JH acquired performed all experiments related to figures 1, 2, 5, 6 and 7, contributed to experiments in figures 3 and 4, and performed all coding and data analysis. RAPV contributed to experiments related to figure 4. AS performed FISH experiments related to figure 3. RBP provided input on methodology and data interpretation. JH and JJL wrote the manuscript after discussions among all authors. JJL supervised the project.

## Acknowledgements

We thank all members of the Letzkus lab, M. Bartos and H. Sprekeler for discussions; A. Wrana and U. Thirimanna for technical assistance; K. Deisseroth and E. S. Boyden, for generously sharing reagents. This work was supported by the German Research Foundation (LE 3804/3-1, LE 3804/4-1, LE 3804/7-1 and LE 3804/8-1 to J.J.L.), and the BrainLinks-BrainTools – IMBIT center (to J.H.), a Peter and Traudl Engelhorn Foundation Postdoctoral Fellowship (to A.S.) and the Brain and Behavior Research Foundation (31041 to A.S.).

## Supplementary information

**Figure S1.**
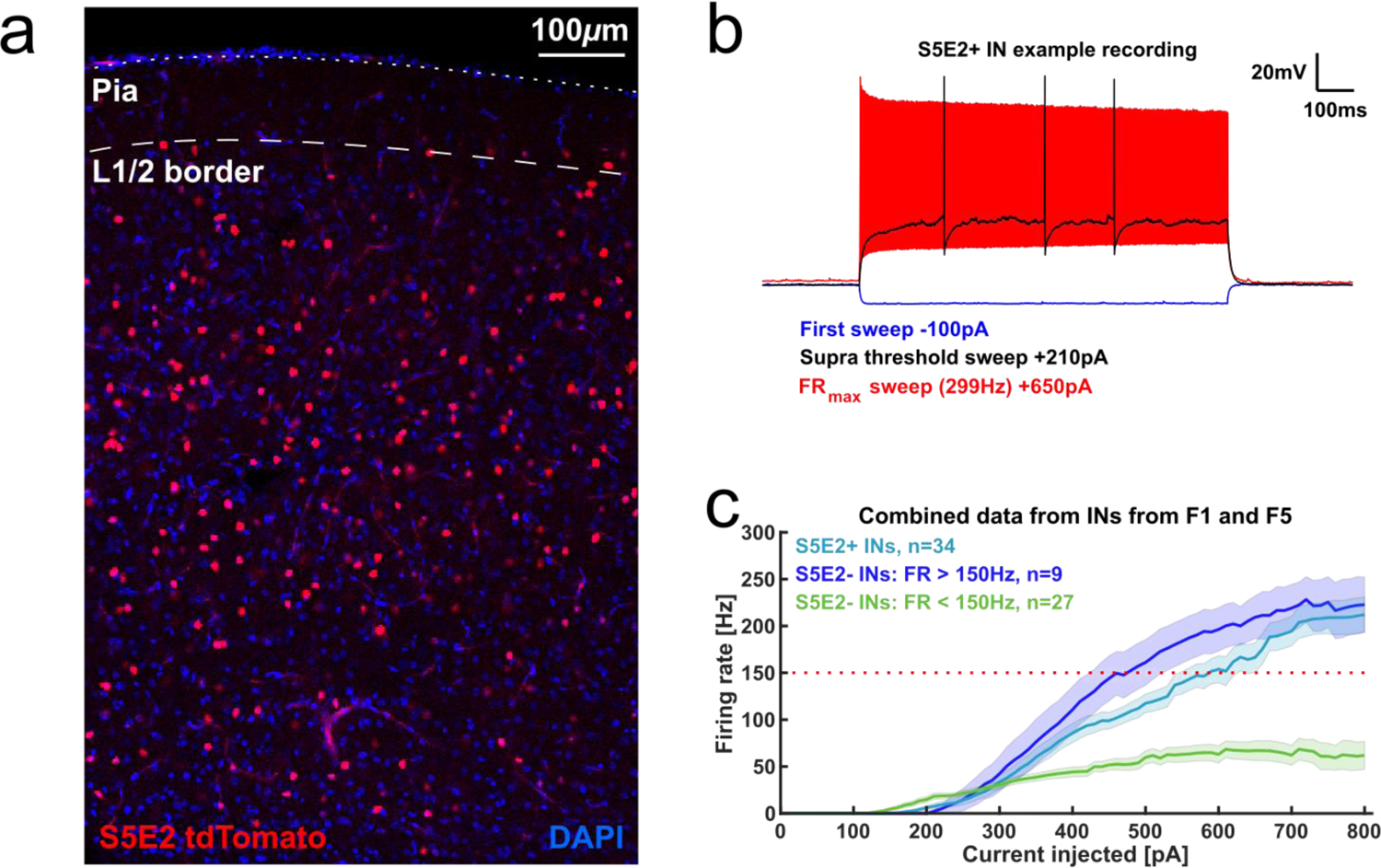
S5E2 labels cortical fast-spiking PV INs. a) Example histology of pAAV-S5E2-Gq-P2A-dTomato. tdTomato expression is absent from cortical L1, consistent with the distribution of PV expression in mouse auditory cortex^13^. Single confocal plane at 20x magnification. b) Example firing pattern of a L2/3 PVIN identified by S5E2 expression. c) I-O plot of PV and non-FSIN sustained firing rates in response to a 1-second current injection. The plot includes data from all INs recorded in F1 and F5 except for genetically identified VIPINs.

**Figure S2.**
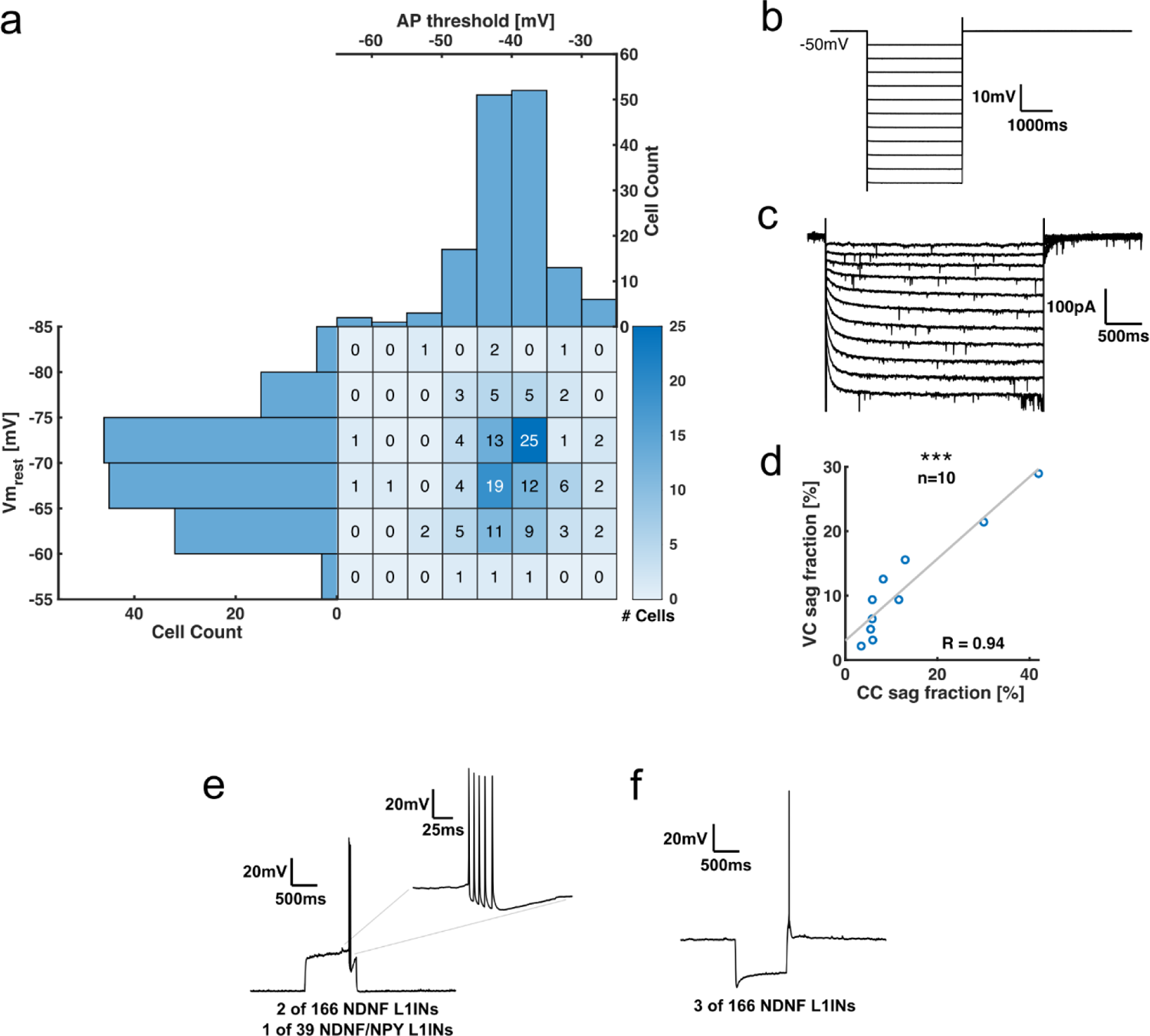
Quality control of electrophysiological recordings in NDNF L1INs. **a)** Marginal distribution of NDNF L1IN AP threshold and resting membrane potential. **b)** Protocol for measuring sag currents in voltage clamp. Cells were clamped to −50mV and hyperpolarized with voltage steps increasing by −5mV each sweep, up to a maximum hyperpolarization of −105mV. **c)** Example recording of the voltage clamp sag fraction protocol **(d)** in a NDNF L1IN. **d)** Correlation of sag fractions measured in voltage clamp and current clamp. Pearson correlation coefficient R=0.94, p=4.1e-5. **e)** Burst firing upon spike onset was rarely observed in late-spiking NDNF or NDNF/NPY L1INs. These cells were excluded from analysis of intrinsic properties. **f)** Rebound spikes were observed rarely after hyperpolarization of NDNF L1INs with large voltage sag.

**Figure S3.**
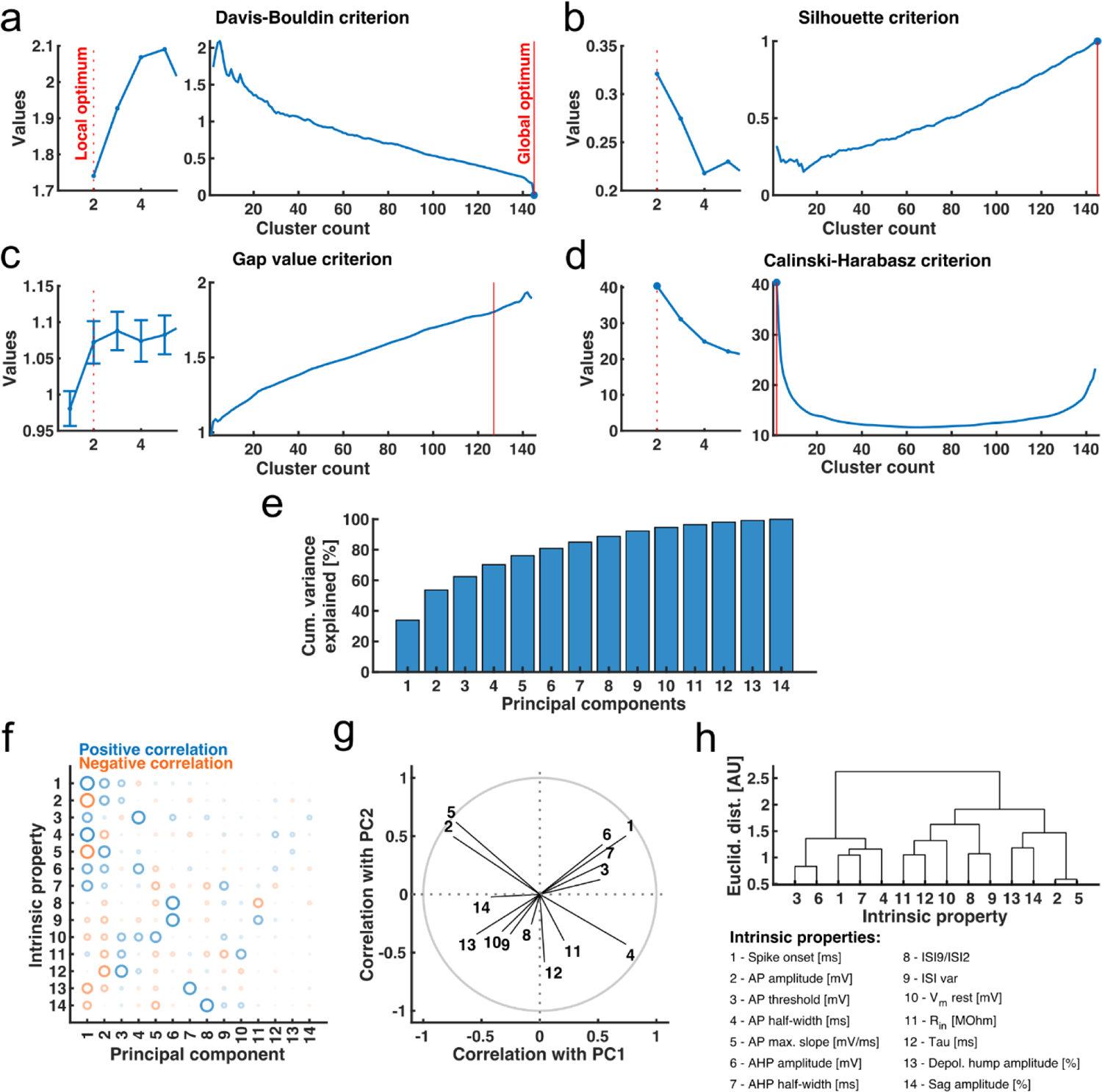
Cluster validation and relation to intrinsic electrophysiological properties. **a-d)** Validation of Ward’s unsupervised clustering using Davis-Bouldin **(a)**, Silhouette **(b)**, Gap value **(c)**, and Calinski-Harabasz **(d)** criteria. Left plots focus on 1 to 5 clusters, dotted red line indicate local optima. Right plots show full distributions, solid red lines indicate global optima. All 4 criteria identify 2 clusters as local optimum number, whereas global maxima vary. **e)** Cumulative variance explained by PCs calculated from NDNF L1IN intrinsic properties. **f)** Contribution of intrinsic properties to PCs measured by their correlation of intrinsic properties with PCs. Circle size indicates correlation strength, circle colors indicate direction of correlation. Absolute correlation strength indicates amount of contribution of a property to a PC. **g)** Correlation of intrinsic properties with first two PCs. Grey unity circle indicates maximum correlation. Magnitude of vectors indicates contribution of a property to first two PCs and thus relevance for clustering. Vectors pointing into similar directions indicate that properties typically occur positively correlated within a given NDNF L1IN, vectors pointing into opposite directions indicate that properties typically occur anticorrelated. **h)** Wards unsupervised clustering of correlation coefficients between all intrinsic properties and all PCs. Short distances between intrinsic properties indicate that they typically occur correlated in a given NDNF L1IN, long distances that they typically occur anticorrelated.

**Figure S4.**
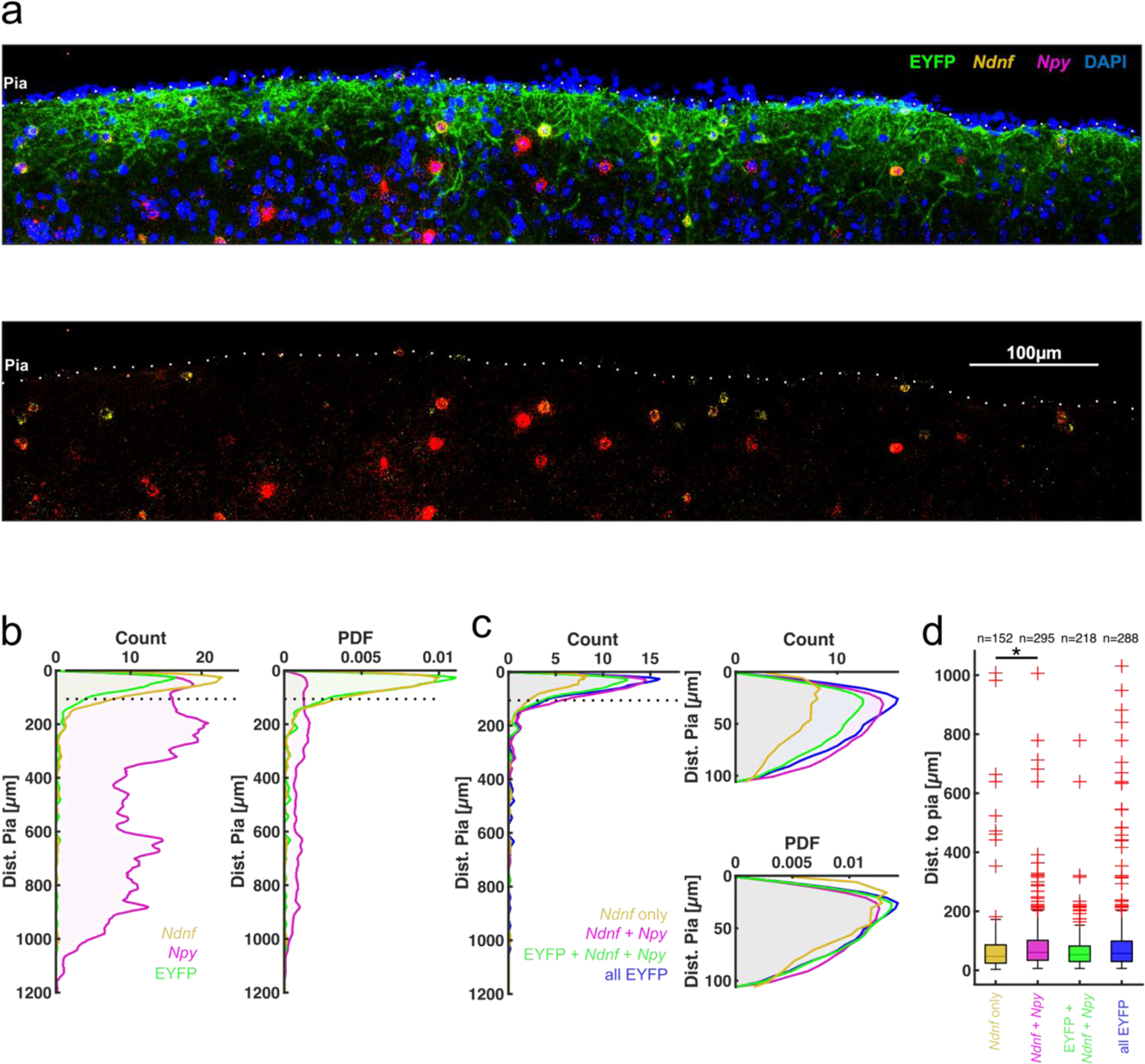
Intersectional genetic targeting of NDNF/NPY L1INs using Boolean logic recombinase approach. **a)** Larger example histology of the validation experiment. **b)** Depth profile of used makers, bin width = 5µm. Yellow: all *Ndnf* expressing cells, Magenta: all *Npy* expressing cells, green: all EYFP expressing cells **c)** Depth profile of combinations of interest for validation, bin width = 5µm. Yellow: cells labeled only for *Ndnf* mRNA, magenta: cells labeled for *Ndnf* and *Npy* mRNA, green: cell labeled for both mRNAs and expressing EYFP, blue: all cells expressing EYFP. **d)** Statistical quantification of c. Kruskal-Wallis test with post-hoc Dunn’s procedure: p_group_=0.015. p*_Ndnf-Ndnf/Npy_*=0.016, p*_Ndnf-_*_EYFP_=0.85, p*_Ndnf_*_-allEYFP_=0.15, p*_Ndnf/Npy_*_-EYFP_=0.22, p*_Ndnf/Npy_*_-allEYFP_=0.92, p_EYFP-allEYFP_=0.81. We observe a slight difference in cortical depth between NPY+ and NPY-NDFN L1INs, but do not find a similar difference between the electrophysiologically defined NDNF L1IN clusters (Fig. 2i).

**Figure S5.**
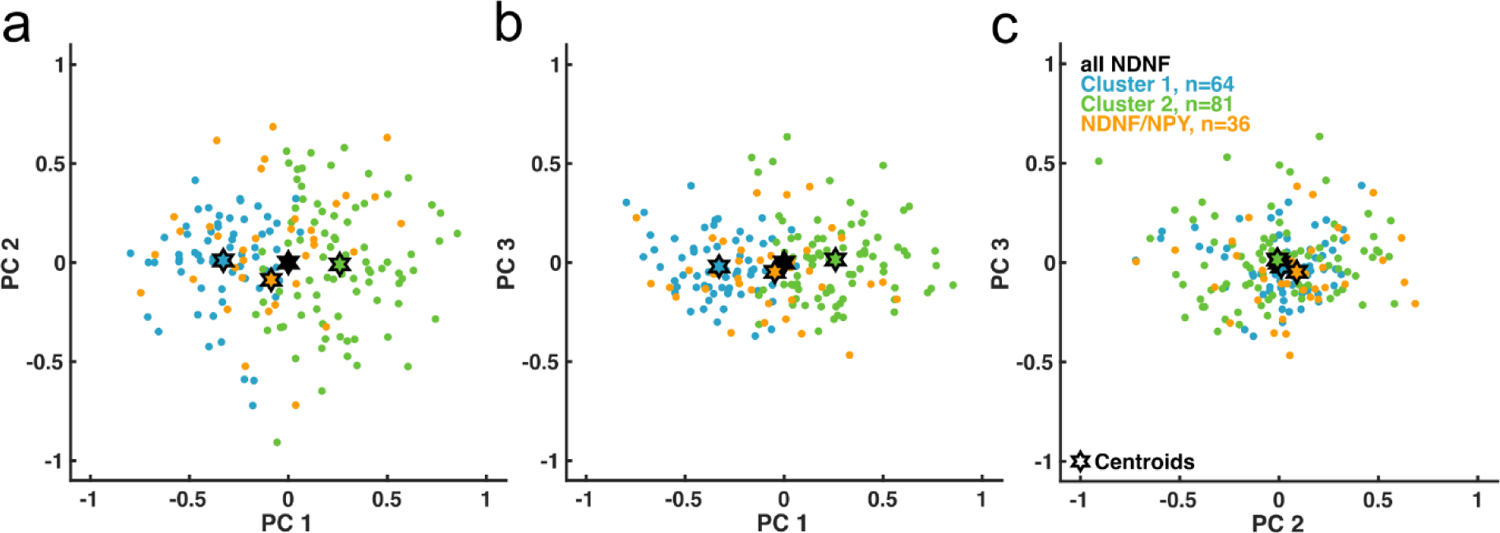
NDNF/NPY L1INs in NDNF L1IN principal component space. a) – c) Projection of NDNF/NPY L1INs into NDNF L1IN principal component space based on NDNF/NPY L1IN intrinsic properties and NDNF L1IN PCA coefficients.

**Figure S6.**
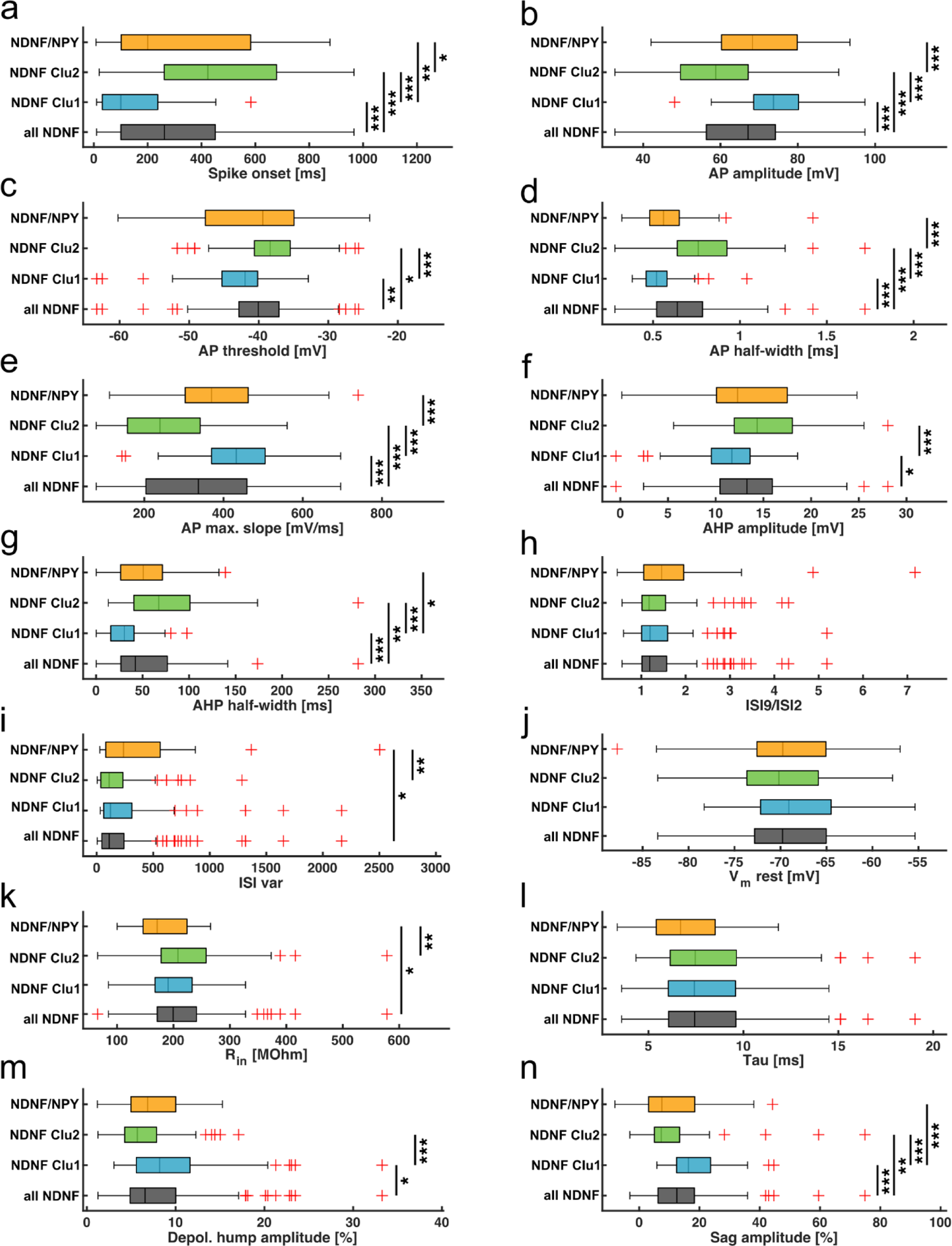
Intrinsic properties by genetic and cluster identity. All statistics done by applying the Kruskal-Wallis-test and post-hoc Dunn’s correction for multiple testing. 1 – all NDNF L1INs, 2 – NDNF Clu1, 3 – NDNF Clu2, 4 – NDNF/NPY INs. a) Spike onset. P_group_=1.1e-12, p_12_=1.1e-5, p_13_=2.8e-4, p_14_=1, p_23_=1.2e-13, p_24_=0.0098, p_34_=0.011. b) Action potential amplitude. P_group_=3.1e-5, p_12_=3.1e-5, p_13_=5.9e-4, p_14_=0.81, p_23_=1.5e-12, p_24_=0.14, p_34_=9.2e-4. c) Action potential threshold. P_group_=7.8e-7, p_12_=0.003, p_13_=0.018, p_14_=0.81, p_23_=1.4e-7, p_24_=0.11, p_34_=0.15. d) Action potential half-width. P_group_=9.5e-14, p_12_=1.5e-5, p_13_=3.5e-4, p_14_=0.083, p_23_=2.6e-13, p_24_=0.79, p_34_=2.5e-6. e) Maximum action potential slope. P_group_=2.1e-12, p_12_=2.6e-5, p_13_=5.1e-4, p_14_=0.48, p_23_=9.5e-13, p_24_=0.33, p_34_=1.4e-4. f) After-hyperpolarization amplitude. P_group_ 2.9e-5, p_12_=0.015, p_13_=0.057, p_14_=1, p_23_=7.1e-6, p_24_=0.25, p_34_=0.25. g) After-hyperpolarization half-width. P_group_=4.7e-10, p_12_=1.3e-4, p_13_=0.0017, p_14_=1, p_23_=5.8e-11, p_24_=0.014, p_34_=0.067. h) Inter-spike interval 9 vs. 2 ratio. P_group_=0.58. i) Inter-spike interval variance. P_group_=0.0028, p_12_=0.67, p_13_=0.81, p_14_=0.023, p_23_=0.15, p_24_=0.51, p_34_=0.0029. j) Resting membrane potential. P_group_=0.58. k) Input resistance. P_group_=0.0078, p_12_=0.84, p_13_=0.92, p_14_=0.031, p_23_=0.37, p_24_=0.43, p_34_=0.0067. l) Membrane time constant. P_group_=0.18. m) Depolarizing hump amplitude. P_group_=4e-4, p_12_=0.047, p_13_=0.13, p_14_=1, p_23_=1.2e-4, p_24_=0.4, p_34=_0.4. n) Voltage sag amplitude. P_group_=9e-9, p_12_=8.9e-4, p_13_=0.0071, p_14_=0.42, p_23_=6.7e-9, p_24_=1.2e-4, p_34_=0.99.

**Figure S7.**
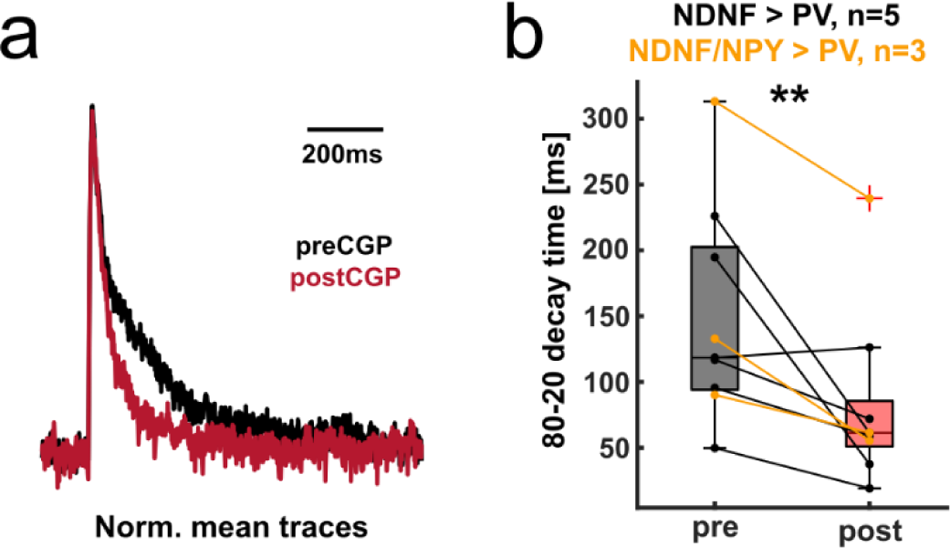
Inhibition of PVINs has as GABA-B receptor component. **a)** Averaged normalized example traces. **b)** c) Effect of CGP on 80-20 decay times in L2/3 PV INs. Data for NDNF L1IN to PVIN and NDNF/NPY L1IN to PVIN connectivity was analyzed together (paired t-test p=0.0089).

**Figure S8.**
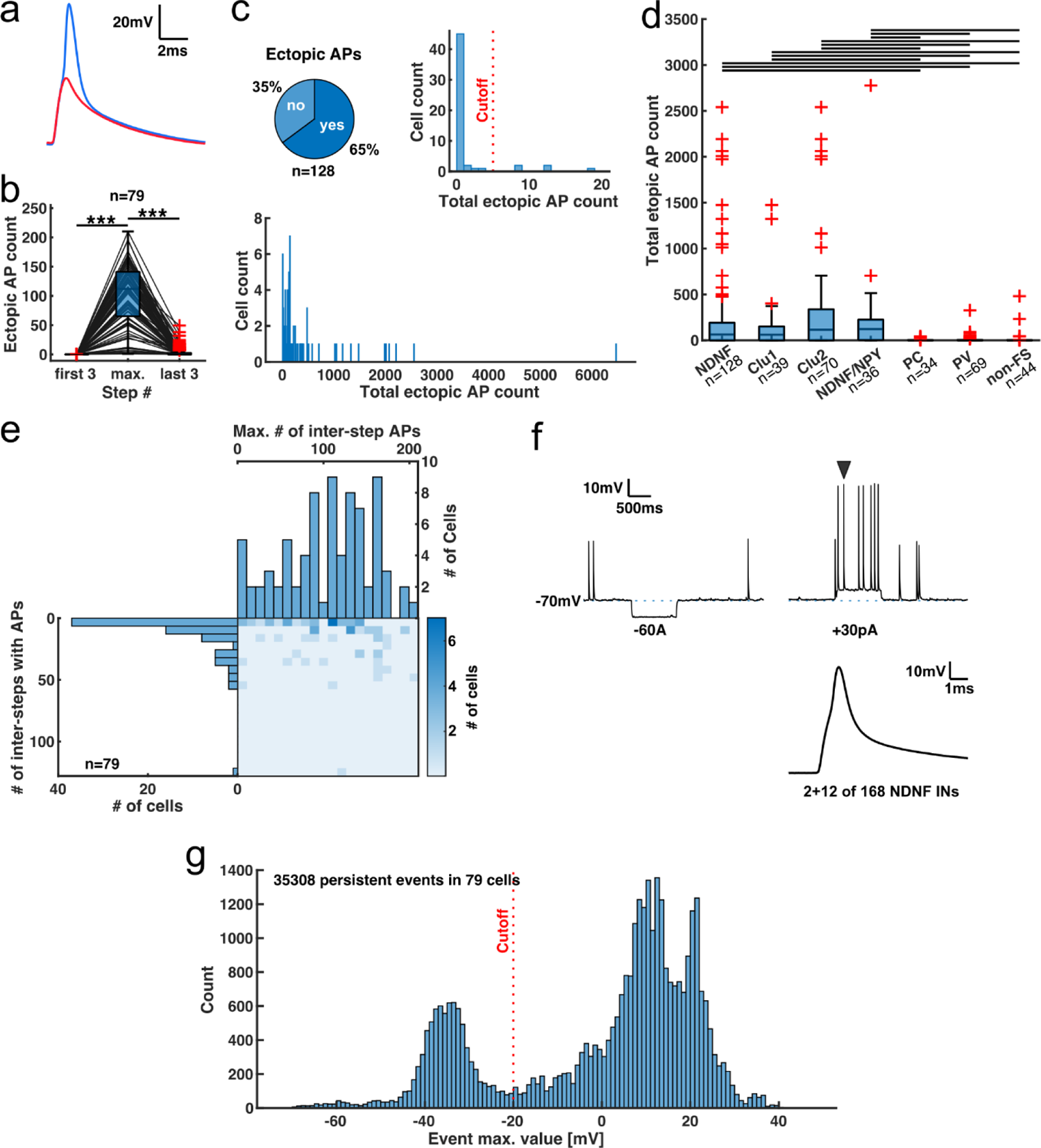
Occurrence of persistent firing. **a)** Overlay of persistent firing spike (blue) and spikelet (red), aligned by action potential crossing time. Persistent firing spikes seem to emerge from persistent firing spikelets. Same data as in F6 d and e. **b)** Boxplot of number of ectopic events, during the first three inter-step intervals with positive current injections, the inter-step interval with the maximum count of ectopic action potentials, and the last three inter-step intervals in each recording. Black lines indicated single data points. Friedman test p_group_=2.8e-32 and post-hoc Dunn’s procedure p_first-max_=0, p_first-last_=0.071, p_max-last_=0. **c)** Upper left: Proportion of NDNF L1INs that show any ectopic action potentials at all. Upper right: Histogram of total ectopic event count, focused on the first 20 bins (bin width = 1). Red line indicates the cutoff chosen for classifying cells as displaying persistent firing. Lower: Full distribution of total ectopic event count in NDNF L1INs (bin width = 10, cells without ectopic events excluded for display clarity). **d)** Total ectopic event count by cell type. Kruskal-Wallis-Test p_group_=4.3e-16. Post-hoc Dunn’s procedure p_NDNF-Clu1_=1, p_NDNF-Clu2_=1, p_NDNF-NDNF/NPY_=1, p_NDNF-PC_=7.3e-6, p_NDNF-PV_=1.5e-6, p_NDNF-nonFS_=1e-5, p_Clu1-Clu2_=1, p_Clu1-NDNF/NPY_=0.98, p_Clu1-PC_=0.003, p_Clu1-PV_=0.0076, p_Clu1-nonFS_=0.0071, p_Clu2-NDNF/NPY_=1, p_Clu2-PC_=1.8e-6, p_Clu2-PV_=6.1e-7, p_Clu2-nonFS_=2.8e-6, p_NDNF/NPY-PC_=8.3e-6, p_NDNF/NPY-PV_=1e-5, p_NDNF/NPY-nonFS_=1.7e-5, p_PC-PV_=1, p_PC-nonFS_=1, p_PV-nonFS_=1. **e)** Marginal distribution of NDNF L1IN number of inter-step intervals with ectopic action potentials and maximum number of action potentials in an inter-step interval. **f)** Example recording of a NDNF L1IN with persistent activity before onset of current-evoked spiking. Upper: Two example sweeps from the same IN with persistent firing spikes and spikelets. Black arrow indicates the action potential on the lower panel with characteristic shoulder. 14 NDNF L1INs showed persistent activity before onset of evoked action potentials. For 12 out of 14, these were typically single spikes. For two neurons, including this example, there was ongoing activity consisting of many events over several sweeps. The latter two cells were excluded from further analysis. **g)** Distribution of maximum voltages reached by ectopic action potentials. There are two clear peaks corresponding to full pF APs and spikelets, respectively. The continuity of the distribution between APs and spikelets corresponds to the fact that values vary between cells, and in particular that pF APs do not always reach the same voltages as current evoked APs (see also **Fig. 7e** for an example). We chose the voltage at the minimum count between the two peaks (−20mV) as a cutoff to distinguish between pF APs and spikelets, indicated in the plot by the red line.

**Figure S9.**
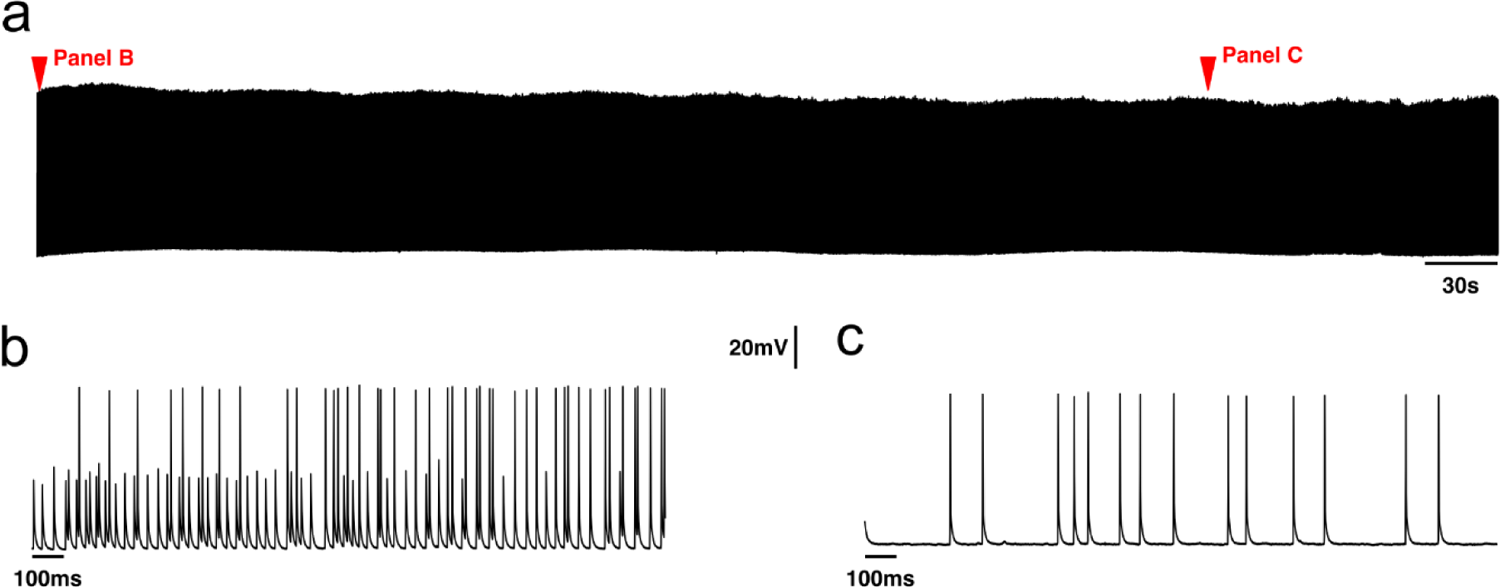
Persistent firing can be very long-lasting. a) Whole recording. b) and c) Focus on beginning (b) and end (c) of recording. As observed during shorter pF episodes during current injection protocols, spikelets tend to be more prevalent at the beginning of the episode.

**Figure S10.**
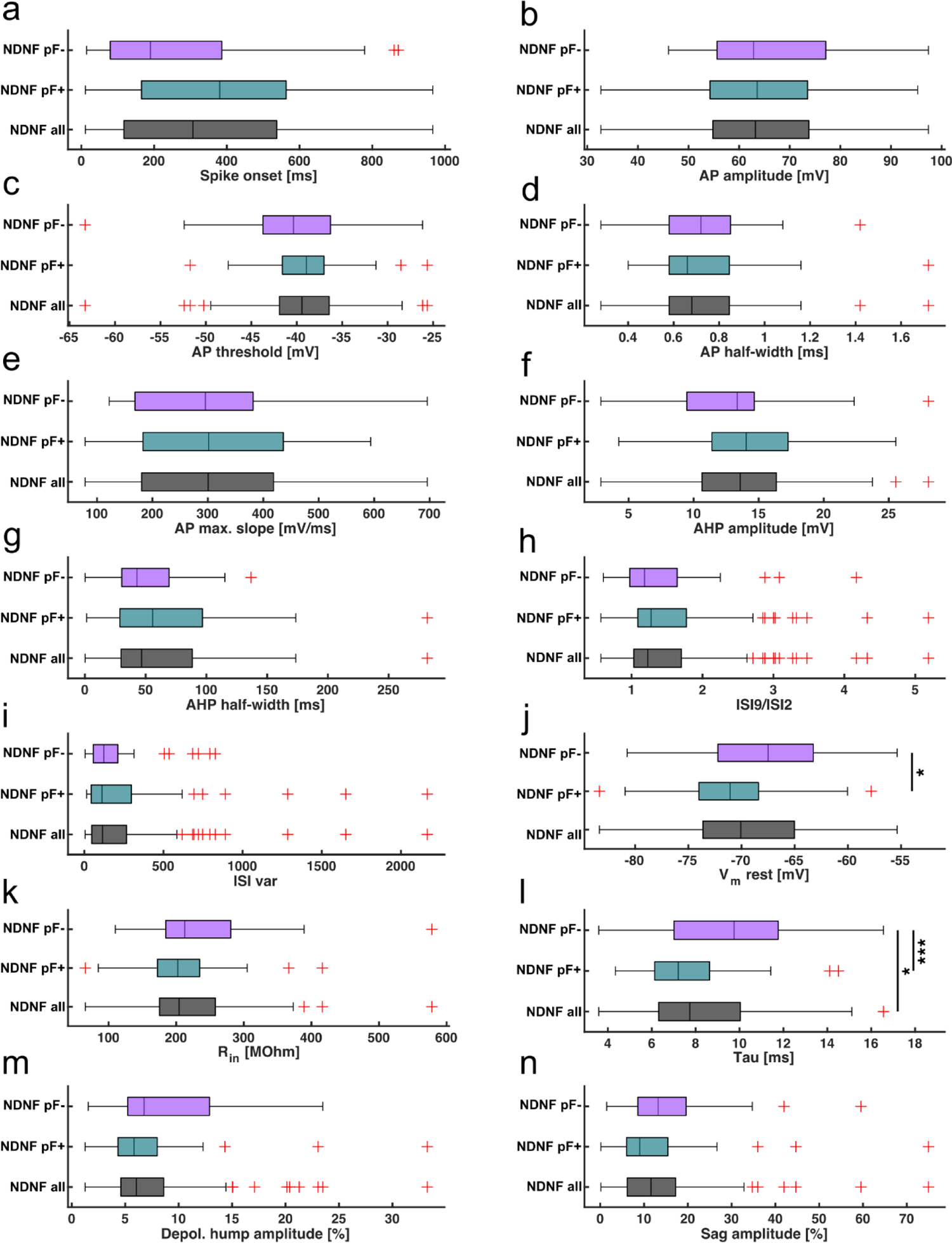
Intrinsic properties of NDNF L1INs by occurrence of persistent firing. All statistics calculated by applying the Kruskal-Wallis-test and post-hoc Dunn’s correction for multiple testing. 1 – all NDNF L1INs, 2 – NDNF pF+ L1INs, 3 – NDNF pF-L1INs. a) Spike onset. P_group_=0.052. b) Action potential amplitude. P_group_=0.77. c) Action potential threshold. P_group_=0.85. d) Action potential half-width. P_group_=1. e) Maximum action potential slope. P_group_=0.96. f) After-hyperpolarization amplitude. P_group_=0.19. g) After-hyperpolarization half-width. P_group_=0.56. h) Inter-spike interval 9 vs. 2 ratio. P_group_=0.56. i) Inter-spike interval variance. P_group_=0.96. j) Resting membrane potential. P_group_=0.024, p_12_=0.48, p_13_=0.18, p_23_=0.019, k) Input resistance. P_group_=0.43. l) Membrane time constant. P_group_=7.3e-4, p_12_=0.2, p_13_=0.029, p_23_=4.3e-4. m) Depolarizing hump amplitude. P_group_=0.12. n) Voltage sag amplitude. P_group_=0.088.

**Figure S11.**
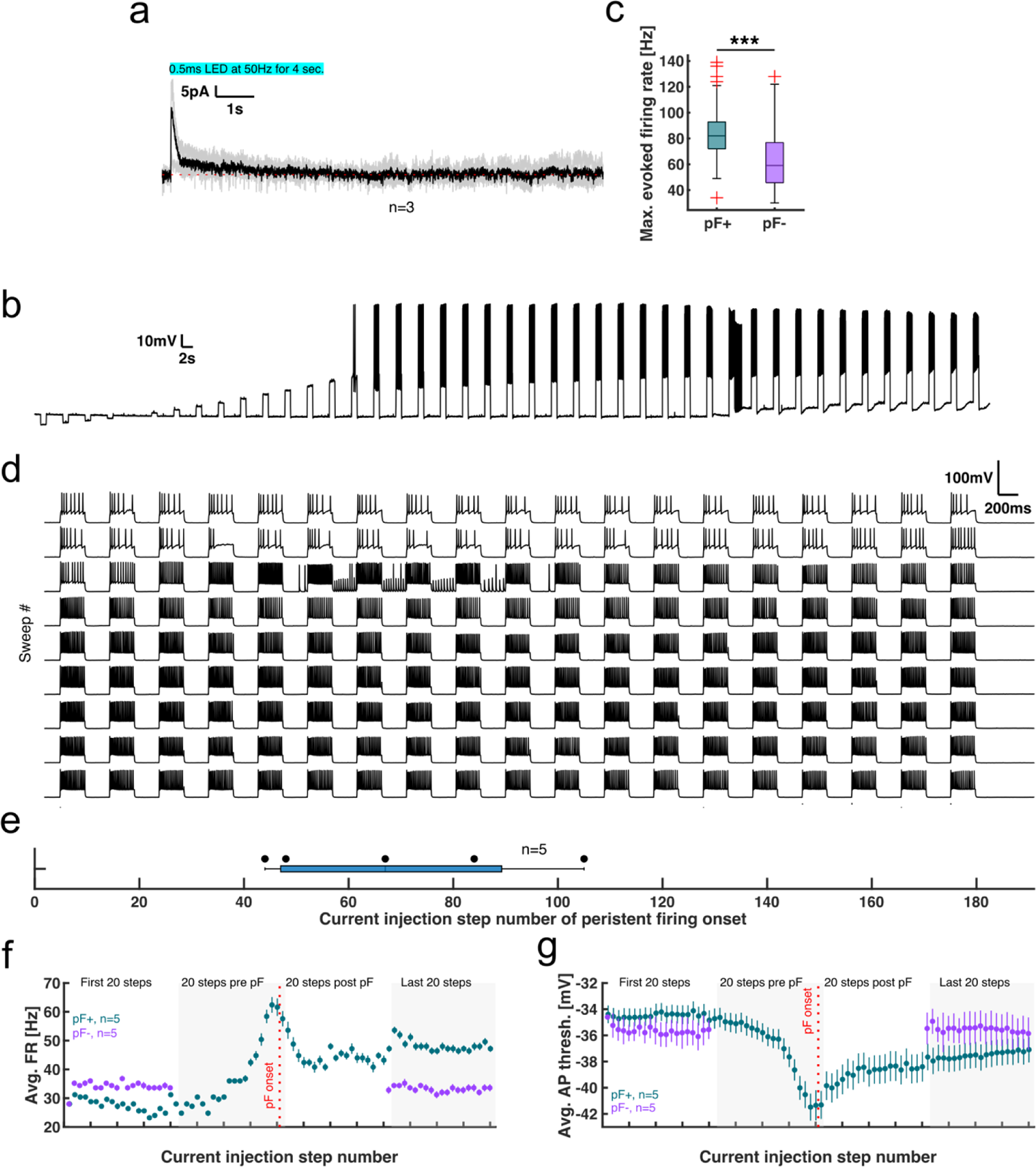
Persistent firing is associated with a state transition in NDNF L1INs. **a)** Complete recording length of data shown in **Fig. 7c**. **b)** Complete recording of recording shown in **Fig. 7e**. **c)** Comparison of maximum current-evoked firing in Ndnf L1INs that do or do not display persistent firing. Wilcoxon rank sum test p=3e-7. **d)** Complete recording of recording shown in **Fig. 7i**. **e)** Onset step number of persistent firing in pF+ NDNF L1INs using the protocol shown in **Fig. 7h**. As pF occurrence is strongly biased towards the first half of the protocol, the absence of pF in the pF-INs is unlikely to be explained by insufficient stimulation. **f)** and **g)** Extended view of current evoked firing rates in **Fig. 7l** (**f**) and action potential thresholds in **Fig. 7n** (**g**). In NDNF L1INs which display persistent firing, firing rates and action potential thresholds stay relatively constant at the beginning of the recording. Towards persistent firing onset firing rates rise first slowly and then sharply and peak shortly before persistent firing onset. After the persistent firing period is over, current evoked firing rates fall again, but stay elevated as compared to start of the recording. Similar dynamics are observed for evoked action potential thresholds, but neither evoked firing rates nor action potential thresholds show any dynamics in NDNF L1INs that do not show persistent firing.

---

**Table S1.**
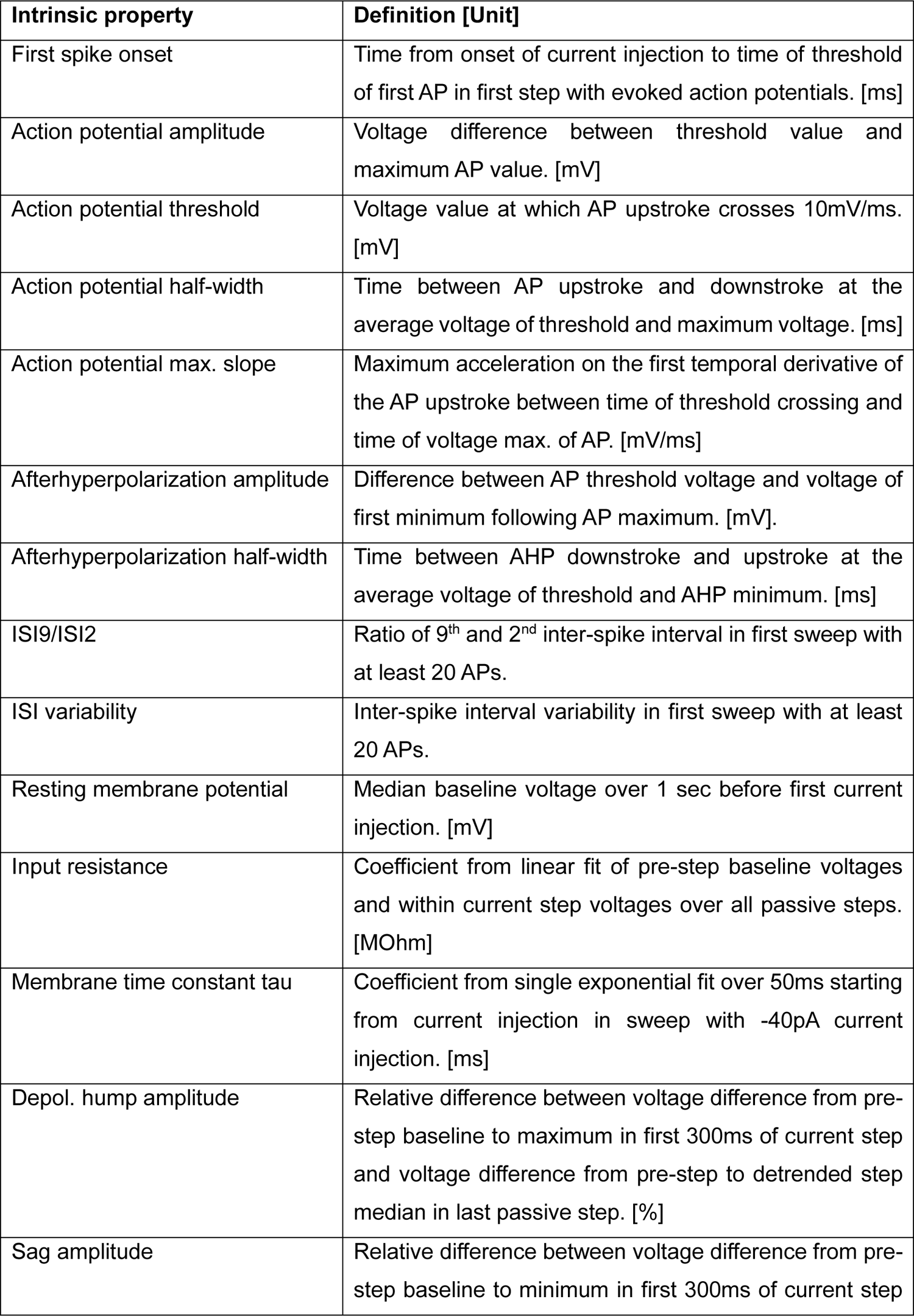
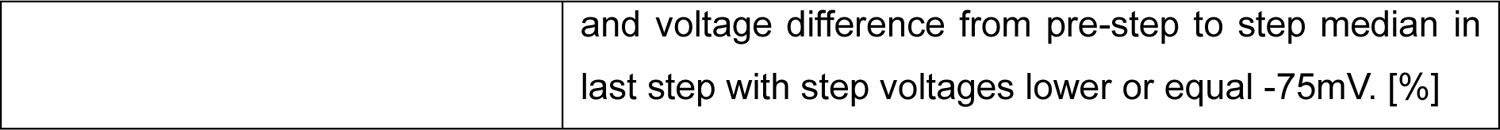
Definition of intrinsic electrophysiological properties. AP and AHP properties (properties 2-7 in the table) were measured in first step with current evoked action potentials and averaged across all APs in that step.

